# MT-MAG: Accurate and interpretable machine learning for complete or partial taxonomic assignments of metagenome-assembled genomes

**DOI:** 10.1101/2022.01.12.475159

**Authors:** Wanxin Li, Lila Kari, Yaoliang Yu, Laura A. Hug

**Affiliations:** School of Computer Science, University of Waterloo, Waterloo, Ontario, Canada; Department of Biology, University of Waterloo, Waterloo, Ontario, Canada

## Abstract

We propose MT-MAG, a novel machine learning-based software tool for the complete or partial hierarchically-structured taxonomic classification of metagenome-assembled genomes (MAGs). MT-MAG is alignment-free, with *k*-mer frequencies being the only feature used to distinguish a DNA sequence from another (herein *k* = 7). MT-MAG is capable of classifying large and diverse metagenomic datasets: a total of 245.68 Gbp in the training sets, and 9.6 Gbp in the test sets analyzed in this study. In addition to complete classifications, MT-MAG offers a “partial classification” option, whereby a classification at a higher taxonomic level is provided for MAGs that cannot be classified to the Species level. MT-MAG outputs complete or partial classification paths, and interpretable numerical classification confidences of its classifications, at all taxonomic ranks. To assess the performance of MT-MAG, we define a “weighted classification accuracy,” with a weighting scheme reflecting the fact that partial classifications at different ranks are not equally informative. For the two benchmarking datasets analyzed (genomes from human gut microbiome species, and bacterial and archaeal genomes assembled from cow rumen metagenomic sequences), MT-MAG achieves an average of 87.32% in weighted classification accuracy. At the Species level, MT-MAG outperforms DeepMicrobes, the only other comparable software tool, by an average of 34.79% in weighted classification accuracy. In addition, MT-MAG is able to completely classify an average of 67.70% of the sequences at the Species level, compared with DeepMicrobes which only classifies 47.45%. Moreover, MT-MAG provides additional information for sequences that it could not classify at the Species level, resulting in the partial or complete classification of 95.13%, of the genomes in the datasets analyzed. Lastly, unlike other taxonomic assignment tools (e.g., GDTB-Tk), MT-MAG is an alignment-free and genetic marker-free tool, able to provide additional bioinformatics analysis to confirm existing or tentative taxonomic assignments.

## Introduction

Metagenome assembled genomes (MAGs) are a technological innovation that has allowed detailed insights into environmental microbial communities, and has strengthened understanding of the uncultured majority of microorganisms ([1], [2]). Accurate taxonomic assignment for these environmentally-derived genomes is a necessary step for identifying populations, making connections across communities and environments, and anchoring hypotheses on metabolic function and roles in biogeochemical cycles ([3], [4]).

As methods for determining phylogeny, evolutionary relationships, and taxonomy, evolved from physical to molecular characteristics, so did many species definitions change. Recently, microbial taxonomy underwent drastic changes through the Genome Taxonomy Database (GTDB, http://gtdb.ecogenomic.org/) in an effort to ensure that taxonomic classifications were standardized, normalized, and evolutionary consistent. In the first GTDB release (i.e., GTDB release 80) nearly 58% of the approximately 84,000 genomes with an attached National Center for Biotechnology Information (NCBI) taxonomy saw a difference in nomenclature above Species-level [5]. With the fourth release (i.e., GTDB release 89) of GTDB, over 30% of the nearly 114,000 genomes with an NCBI taxonomy (out of 143,000 total genomes in GTDB at the time) saw a change in the assigned Species taxon [6].

In the absence of a definitive ground truth, any existing and newly proposed Species clusters would benefit from additional bioinformatics analysis by complementary genome-based classification methods, to confirm tentative taxonomic assignments. Even though existing taxonomic assignment tools (e.g., Kraken 2 [7], BERTax [8],

GTDB-Tk [9]) have achieved good classification accuracies on benchmarked tasks, they are constrained by various limitations, as described below.

Alignment-based tools (e.g., GTDB-Tk [9]) require DNA sequences to be aligned to reference sequences to obtain sequence similarities [10]. In addition, alignment-based tools assume that homologous sequences are composed of a series of linearly arranged and more or less conserved sequence stretches, assumptions that may not always hold due to high mutation rates, frequent genetic recombination events, etc. [11]. Lastly, the utility of alignment-based tools is limited by their often prohibitive consumption of runtime and computational memory.

Genetic marker-based tools such as IDTAXA [3] and GTDB-Tk [9] rely on taxonomic markers (e.g., 16S ribosomal RNA genes, internal transcribed spacers) to identify microorganisms. The use of genetic marker-based tools is limited by the fact that partial genomes frequently lack major markers. The absence of major markers could be caused, e.g., by the genome not being sequenced to a sufficient depth to assemble well, resulting in markers of interest possibly missing from the assembly [12]. An additional reason could be that fragments carrying the markers do not bin with the rest of the genome, which is a frequent problem with 16S ribosomal RNA genes [13].

At the other end of the spectrum, alignment-free tools based on *k*-mer frequencies (e.g., DeepMicrobes [14], CLARK [15]) do not rely on alignment or genetic markers, and instead use *k*-mer frequencies as the input feature. However, existing *k*-mer-based tools are also limited by, e.g., the fact that they are only capable of taxonomic assignment at specific taxonomic levels (e.g., Genus, Species), and a lack of interpretability of their predicted taxonomic assignments.

To address these limitations, we propose MT-MAG, a **m**achine learning-based **t**axonomic assignment tool for **m**etagenome-**a**ssembled **g**enomes. Unlike most other tools (e.g., GTDB-Tk) MT-MAG is an *alignment-free* and *genetic marker-free* software tool. MT-MAG uses only their *k*-mer frequencies to distinguish DNA sequences from one another (for the datasets in this paper, the optimal value of *k* was empirically determined to be *k* = 7). In addition, by using a hierarchical local classification approach MT-MAG is able to provide partial classifications (at higher taxonomic levels than, e.g., Species) for MAGs that it cannot confidently classify at the Species level. Lastly, for a query genome, MT-MAG outputs not only a classification path, but also a numerical classification confidence of its prediction, at each taxonomic rank. The main contributions of this paper are:

- **Partial Classification:** A feature of MT-MAG is that it outputs partial classifications for the majority of sequences that it cannot confidently classify at the Species level. This results in an average of 95.13% of the genomes in the datasets analyzed being either partially or completely classified. In particular, MT-MAG completely classifies, on average, 88.84% of the test sequences to the Phylum level, 88.39% to the Class level, 86.81% to the Order level, 81.17% to the Family level, and 71.13% to the Genus level.
- **Interpretability:** MT-MAG outputs numerical classification confidences for its classifications, at all taxonomic ranks along the classification path.
- **Weighted Classification Accuracy:** To assess the performance of MT-MAG, we introduce the “weighted classification accuracy,” a performance metric defined as the weighted sum of the proportions of complete and partial classifications. To the best of our knowledge, this is the first metric that incorporates a weighting scheme which reflects the fact that partial classifications at different ranks are not equally informative.
- **Large Datasets:** MT-MAG is capable of classifying large and diverse metagenomic datasets. The two datasets analyzed in this paper are: genomes from human gut microbiome species (training set 6.15 Gbp, test set 7.42 Gbp), and bacterial and archaeal genomes assembled from cow rumen metagenomic sequences (training set 239.53 Gbp, test set 2.18 Gbp).
- **Superior Performance:** MT-MAG achieves an average of 87.32% in weighted classification accuracy, for the datasets analyzed. In particular, at the Species level (the only comparable taxonomic rank with DeepMicrobes), MT-MAG outperforms DeepMicrobes by an average of 34.79% in weighted classification accuracy. In addition, MT-MAG is able to completely classify an average of 67.70% of the sequences at the Species level, compared to DeepMicrobes, which only classifies 47.45%.

## Materials, Methods, and Performance Metrics

This section describes the datasets used, outlines the MT-MAG classification algorithm, and defines the performance metrics used to analyze MT-MAG’s performance and to compare it with that of DeepMicrobes.

### Materials: Datasets and task description

Two different tasks were performed in the computational experiments of this study, called **Task 1** and **Task 2**. The dataset analyzed in Task 1 was selected for direct performance comparison purposes, as it was the dataset analyzed by DeepMicrobes [14]. More specifically, the MT-MAG training set in Task 1 was based on representative genomes from species in human gut microbiomes, and the test set comprised high-quality MAGs reconstructed from human gut microbiomes from a European Nucleotide Archive study [16]. The MT-MAG training set in Task 2 was based on representative and non-representative microbial genomes from GTDB r202, and the test set comprised 913 “draft” bacterial and archaeal genomes assembled from rumen metagenomic sequence data derived from 43 Scottish cattle [17].

The rationale behind the selection of DeepMicrobes for a benchmark comparison with MT-MAG is as follows. Like MT-MAG, DeepMicrobes is a machine learning-based alignment-free and genetic marker-free metagenomic taxonomic assignment tool that uses *k*-mer frequencies as input feature to predict taxonomic assignments of short reads at the Genus and Species level. DeepMicrobes has demonstrated better performance at the Species level classification, and better comparative accuracy in Species abundance estimation over other state-of-the-art tools, see [**?**, 7, 15, 18–20]. In addition, like MT-MAG, DeepMicrobes estimates classification confidences: The reads with classification confidences below a (constant) threshold are considered to be *unclassified reads*, while the rest are considered to be *classified reads*. Within the set of classified reads, the reads whose classified Species taxa are the same as their ground-truth Species taxa are considered to be *correctly classified reads*. Lastly, to the best of our knowledge, DeepMicrobes is the only other taxonomic assignment tool that enables probabilistic classification using machine learning classifiers and has attempted to classify large datasets (e.g., thousands of species), similar to MT-MAG’s design goals.

As MT-MAG and DeepMicrobes have different requirements on their inputs, in that MT-MAG ideally requires the training sequences to be *>* 10,000 bp, while DeepMicrobes operates with short reads, the datasets were prepared separately for MT-MAG and DeepMicrobes.

We conclude these general remarks on the datasets used in this study with a discussion on the reference labels that were used for both the training sets and test sets. We first note that the NCBI [22] labels are outdated, due to the lack of consensus on uncultivated taxa naming conventions [23]. In contrast, in GTDB a consistent naming scheme was achieved by naming uncultivated taxa as ‘Genus name’ sp1, ‘Genus name’ sp2, and so on [5]. Second, we note that GTDB provides a complete taxonomic hierarchy with no inconsistencies in naming, based on standardized phylogenetic distances used to define taxonomic ranks [6]. Third, we observe that the numerical labels used by DeepMicrobes are not biologically meaningful, and cannot be extended to other datasets. Consequently, in this study we used as reference labels the labels obtained by running GTDB-Tk [9] (based on GTDB R06-RS202, April 27, 2021), since they represent the current consensus of the scientific community on microbial taxonomy.

### Task 1: Sparse training set

The dataset for Task 1 was specifically chosen so as to allow a direct comparison between the quantitative performance of MT-MAG and that of DeepMicrobes (see “Performance metrics”). Since the genomes that the training sets for Task 1 were based on comprise only 2.4 % of the GTDB at the Species level, in the remainder of the paper this task will be referred to as *Task 1 (sparse)*.

We first note that we were unable to replicate the classification accuracies reported by DeepMicrobes, using the datasets and software provided in [14]. Absent the possibility to reproduce the results in [14] *ab initio*, and to give DeepMicrobes the best possible scenario for comparison, we opted for the alternative of using the already trained Species classification model reported in [14].

The training set for the Species classification model provided by [14], consisted of reads extracted from 2,505 representative genomes of human gut microbial species. These genomes were identified previously by a large-scale assembling study of the species in human gut microbiomes, and are available at the FTP website ftp://ftp.ebi.ac.uk/pub/databases/metagenomics/umgs_analyses. This genome set comprised 1,952 MAGs, and 553 microbial gut Species-level genome representatives from the human-specific reference (HR) database. This 2,505 genome set was referred to in [14] as *“HGR*.*”* Starting from HGR, DeepMicrobes [14] first assigned each species a numerical label from 0 to 2,504 (inclusive). Secondly, using the ART Illumina read simulator [24], 100,000 150 basepair (bp) paired-end reads were simulated from HiSeq 2500 error model with the mean fragment size of 200 and standard deviation of 50 bp per species. Thirdly, the simulated reads were trimmed from the 3′ end to 75–150 bp in equal probability. Lastly, these trimmed simulated read sets with their numerical labels from 0 to 2504 (inclusive) were used as the input to DeepMicrobes. The total size of the training set of this Species classification model trained by DeepMicrobes is 56.03 Gbp.

The test set of DeepMicrobes was prepared in [14] in a similar way to the training set, and it comprised twenty-thousand 75-150 bp trimmed paired-end reads per MAG, simulated using ART Illumina from 3,269 high-quality MAGs reconstructed from human gut microbiomes from a European Nucleotide Archive study titled “A new **g**enomic **b**lueprint of the **h**uman **g**ut **m**icrobiota” (GBHGM) [16]. The reference taxonomic labels for the test set were derived by running GTDB-Tk. The total test set size for DeeMicrobes is 14.71 Gbp.

The training set of MT-MAG was prepared as follows. Since MT-MAG uses an enhanced version of MLDSP as a subprocess (see “Methods: MT-MAG algorithm”), which achieves optimal performance when the input sequence length exceeds 10,000 bp, all contigs in HGR that were shorter than 5,000 bp were discarded. The remaining 14,358 contigs comprised the training set of MT-MAG, totalling 6.15 Gbp. The process by which MT-MAG handles the special case of imbalanced classes, and the special case of the input dataset being too large to be loaded in memory are described in *S1 Appendix: Materials - Datasets and task description* Section 3.

The test set of MT-MAG comprised 3,269 full MAGs in GBHGM. The total size of the test set of MT-MAG is 7.42 Gbp.

Finally, to compare the DeepMicrobes classification results with those of MT-MAG, we post-processed the numerical labels of the reads in the DeepMicrobes training set, as follows. Recall that the reads in the training set were simulated from real genomes in the HGR database. Post-processing the numerical label of a read in the training set entailed using GTDB-Tk to obtain the GTDB reference label of its originating genome, and this GTDB label was then associated to the numerical label of that read.

### Task 2: Dense training set

The training sets used in Task 2 were based on genomes comprising 7.7% of GTDB taxonomy, hence this task will thereafter be referred to as *Task 2 (dense)*.

The training set of MT-MAG was prepared using GTDB R06-RS202. Note that the sizes of the genomes in GTDB are significantly larger than those of genomes in HGR. Most GTDB MAGs contain multiple contigs per genome. All contigs belonging to a given genome were pseudo-concatenated into a single sequence, by adding the symbol “O” between contigs, so as to avoid creating artificial *k*-mers at the junction of contigs. Then, 4 non-overlapping fragments of length 100,000 bp were selected from each such genome, using four random starts. The 4 obtained fragments belonging to the same genome were again pseudo-concatenated to form a *representative genomic fragment* for that genome. To ensure that we had a sufficient number of representative genomic fragments to perform cross-validation, the above sampling process was repeated 20 times for each genome, resulting in 20 separate representative genomic fragments with the same genome label. The total size of the training set of MT-MAG is 239.53 Gbp. The process by which MT-MAG handles the special case of imbalanced classes, and the special case of the input dataset being too large or too small, are described in *S1 Appendix: Materials - Datasets and task description* Section 3.

Regarding the preparation of the training set of DeepMicrobes, we note that the only way to enable the DeepMicrobes model to handle the training stage of Task 2 (dense) (training set comprising 49,871 species) is to increase the size of *k*-mer. However, according to our computational experiments, *k* = 12 is the largest value that can be handled by a 24Gb GPU memory. Thus, to make the benchmarking comparison with MT-MAG possible, we opted to include in the DeepMicrobes training set only reads from genomes whose species are present in the DeepMicrobes test set (601 species). Note that, without drastically trimming its training set this way, DeepMicrobes would have to load into its memory all 49,871 Species in GTDB, the value of *k* would have to be increased to obtain a satisfactory accuracy, and both these factors would lead to DeepMicrobes crashing due to RAM limitations. Note also that this design decision actually gives a competitive advantage to DeepMicrobes in this comparison, as it is likely to increase DeepMicrobes’ classification accuracy by a large amount.

Following this design choice, the training set of DeepMicrobes was prepared from the representative and non-representative genomes of the afore-mentioned 601 species, in a similar way to the training set of DeepMicrobes in Task 1 (sparse). Approximately thirty-thousand 75-150 bp paired-end reads were simulated per species, and each species was assigned a numerical label between 0 and 600 (see *S1 Appendix: Materials -Datasets and task description* Section 4 for details).

The test set of MT-MAG comprised 913 full microbial genomes from metagenomic sequencing of cow rumen, which were derived from 43 Scottish cattle [17]. The total sequence length of the test set of MT-MAG is 2.18 Gbp.

The test set of DeepMicrobes (reads) was prepared from the 913 full microbial genomes [17], in a similar way to the test set of DeepMicrobes in Task 1 (sparse). In the end, 10,000 75-150bp trimmed simulated paired-end reads per MAG were generated as the input to DeepMicrobes.

The total size of the test set of DeepMicrobes is 2.04 Gbp, and 18,143,340 reads were simulated.

Table 1 provides a summary of the total number of basepairs analyzed, number of FASTA files, number of contigs/reads, and the range of contig/read length for training and test sets in Task 1 (sparse) and Task 2 (dense), for MT-MAG and DeepMicrobes.

**Table 1.**
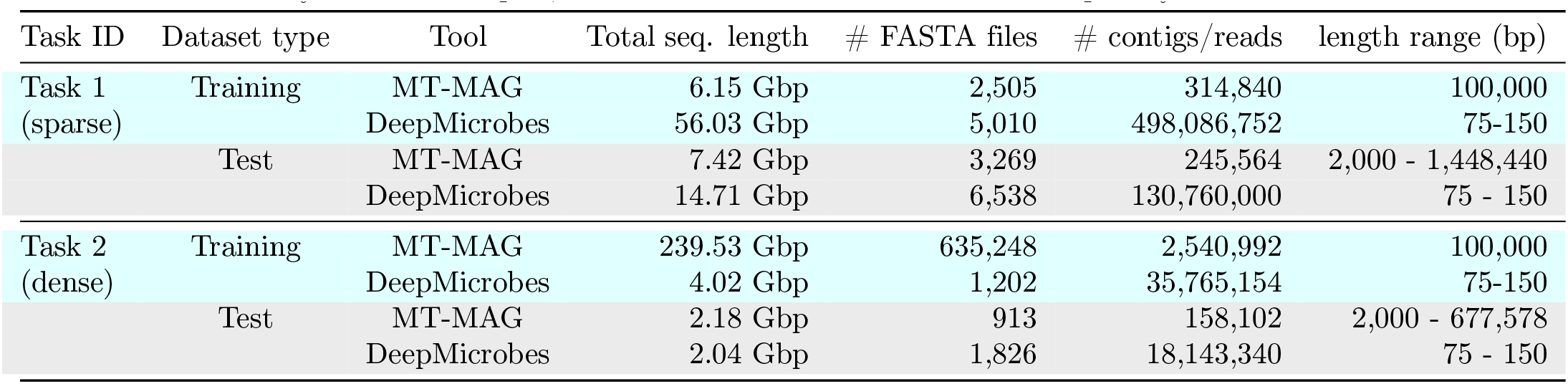
Summary of total number of basepairs analyzed, number of FASTA files, number of contigs/reads, and the range of contig/read lengths for the training and test sets in Task 1 (sparse), and Task 2 (dense) for MT-MAG and DeepMicrobes. Note that contigs come directly from the samples, while reads are simulated from the samples by the ART simulator.

| Task ID | Dataset type | Tool | Total seq. length | # FASTA files | # contigs/reads | length range (bp) |
| --- | --- | --- | --- | --- | --- | --- |
| Task 1<br>(sparse) | Training | MT-MAG | 6.15 Gbp | 2,505 | 314,840 | 100,000 |
|  |  | DeepMicrobes | 56.03 Gbp | 5,010 | 498,086,752 | 75-150 |
|  | Test | MT-MAG | 7.42 Gbp | 3,269 | 245,564 | 2,000 - 1,448,440 |
|  |  | DeepMicrobes | 14.71 Gbp | 6,538 | 130,760,000 | 75 - 150 |
| Task 2<br>(dense) | Training | MT-MAG | 239.53 Gbp | 635,248 | 2,540,992 | 100,000 |
|  |  | DeepMicrobes | 4.02 Gbp | 1,202 | 35,765,154 | 75-150 |
|  | Test | MT-MAG | 2.18 Gbp | 913 | 158,102 | 2,000 - 677,578 |
|  |  | DeepMicrobes | 2.04 Gbp | 1,826 | 18,143,340 | 75 - 150 |

### Methods: MT-MAG algorithm

This section describes the hierarchically-structured local classification approach used by MT-MAG, the eMLDSP subprocess that is at the core of MT-MAG, and the two main phases of MT-MAG (training and classifying).

#### A hierarchically-structured local classification approach

Taxonomic assignment is a problem of hierarchical classification, whereby input items are grouped according to a hierarchy. A hierarchy can be formalized as a directed acyclic graph where every node can be reached by a unique path from the root node (see Fig 1). In machine learning, there are generally three types of approaches to hierarchical classification [25]. The simplest approach is *flat classification* where all parent nodes are ignored, and a single classifier is trained to classify each instance directly into a leaf node. The second approach is the so-called *big bang* classification where a single classifier is trained for all nodes in the hierarchy. The third approach is the *hierarchically-structured local classification*, whereby one multi-class classifier is trained for each parent-to-child relationship. This third approach is an iterative classifying process where instances classified to a child node are then further classified with the next-level classifier, where the child node is now the parent node for the next-level classifier.

**Fig 1.**
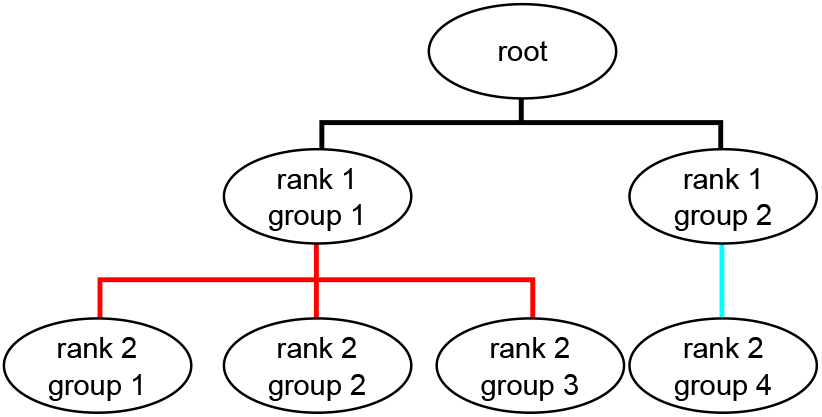
A sample hierarchy (taxonomy) with three parent-to-child relationships. A parent node with all its children nodes forms a parent-to-child relationship. A parent node without a child node is called a leaf node. The level of a node is the length of path from that node to the root node. The part highlighted in red is a multi-child classification, while the part highlighted in cyan is a single-child classification.

In contrast with DeepMicrobes which uses flat classification, MT-MAG uses hierarchically-structured local classification, for reasons detailed below. First, in the case of flat classification, an erroneous classification of a DNA sequence directly at the Species level is more likely, due to the very large number of classes at the Species level. This, in turn, results in a higher likelihood of placing the sequence into an erroneous higher-level taxonomic rank, e.g., Order. Such a serious misplacement is less likely to happen with hierarchically-structured local classification, whereby a sequence passes through multiple classifications, from higher to lower taxonomic ranks, thus providing multiple check-points for the identification of an incorrect classification. For example, an incorrect Order classification could be prevented if any of the classifications prior to and including this level are deemed “uncertain.”

In addition, in the case of flat classification, if the classification confidence of a sequence into a Species taxon does not meet the required confidence level, this sequence is simply deemed “unclassified,” with no further information being provided. In contrast, the hierarchically-structured local classification provides the option of partial classification and can output partial classification paths for such sequences, even if their Species level classification is uncertain.

Finally, flat classification requires significantly more computational time and memory resources, because it involves a single big classification task wherein all the training sequences are loaded into memory simultaneously. In contrast, a hierarchically-structured local classification approach involves multiple smaller classification tasks and, for each classification task, one only needs to load into memory the sequences pertaining to the specific parent taxon being classified at this step in the hierarchy. In particular, for classifications at higher taxonomic ranks, one can use, e.g., only representative genomes as opposed to all of the genomes available for that parent taxon.

#### The enhanced MLDSP (eMLDSP) subprocess

MT-MAG uses an enhanced version of MLDSP, an alignment-free software tool that combines supervised machine learning techniques with digital signal processing for ultrafast, accurate, and scalable genome classification at all taxonomic ranks [26, 27].

The inputs to MLDSP are pseudo-concatenated DNA sequences, together with their reference taxonomic labels. After selecting a value for the parameter *k*, each such input DNA sequence is converted into a numerical vector containing the counts of all of its *k*-mers, where a *k*-mer is defined as a DNA subsequence of length *k* that does not contain the symbol “O” (used during the pseudo-concatenation process), or the symbol “N” (representing an unidentified nucleotide). Each *k*-mer count vector is then converted into a *k-mer frequency vector*, via dividing its *k*-mer counts by the total length of the sequence (excluding “O”s and “N”s). These *k*-mer frequency vectors are computed via order *k* Frequency Chaos Game Representation of a DNA sequence (*FCGR*_*k*_) [28–30], and used as the input to MLDSP.

The optimal value of *k* is in general dependent on the dataset. For this study, we conducted preliminary experiments with values of *k* between 7 and 11, and these values yielded similar accuracy results. Thus, the value *k* = 7 was selected, since it resulted in a high classification accuracy while requiring the least computational resources.

MLDSP consists of two main steps: (a) *Preprocessing*, whereby several different classifiers’ performance is evaluated by 10-fold cross validation, and (b) *Classify*, whereby MLDSP first trains the classifiers using the entire training set *(Classify-Training)*, and then classifies new DNA sequences in the test set *(Classify-Classification)* (see *S2 Appendix: Methods - MT-MAG algorithm* Section 1).

MT-MAG uses an enhanced version of MLDSP, called *eMLDSP* (enhanced MLDSP) as a subprocess. The eMLDSP subprocess augments MLDSP in several significant ways. First, it augments MLDSP by adding the capability to handle the special case where the parent taxon has only one child taxon, as well as by adding the new feature of computing classification confidences for its classifications. Second, it adds a stopping threshold picking algorithm, called “STP algorithm,” which is at the core of the partial classification option feature of MT-MAG. Specifically, the STP algorithm provides an individual stopping threshold for each parent-child pair, at each taxonomic level, as opposed to the one-size-fits-all stopping threshold of DeepMicrobes at the Species level. Third, eMLDSP combines the hierarchically-structured local classification with the result of the STP algorithm to output “uncertain classification,” if the classification confidence is below the stopping threshold.

Fig 2 provides an overview of eMLDSP, including eMLDSP (Preprocessing), eMLDSP (Classify-Training) and eMLDSP (Classify-Classification).

**Fig 2.**
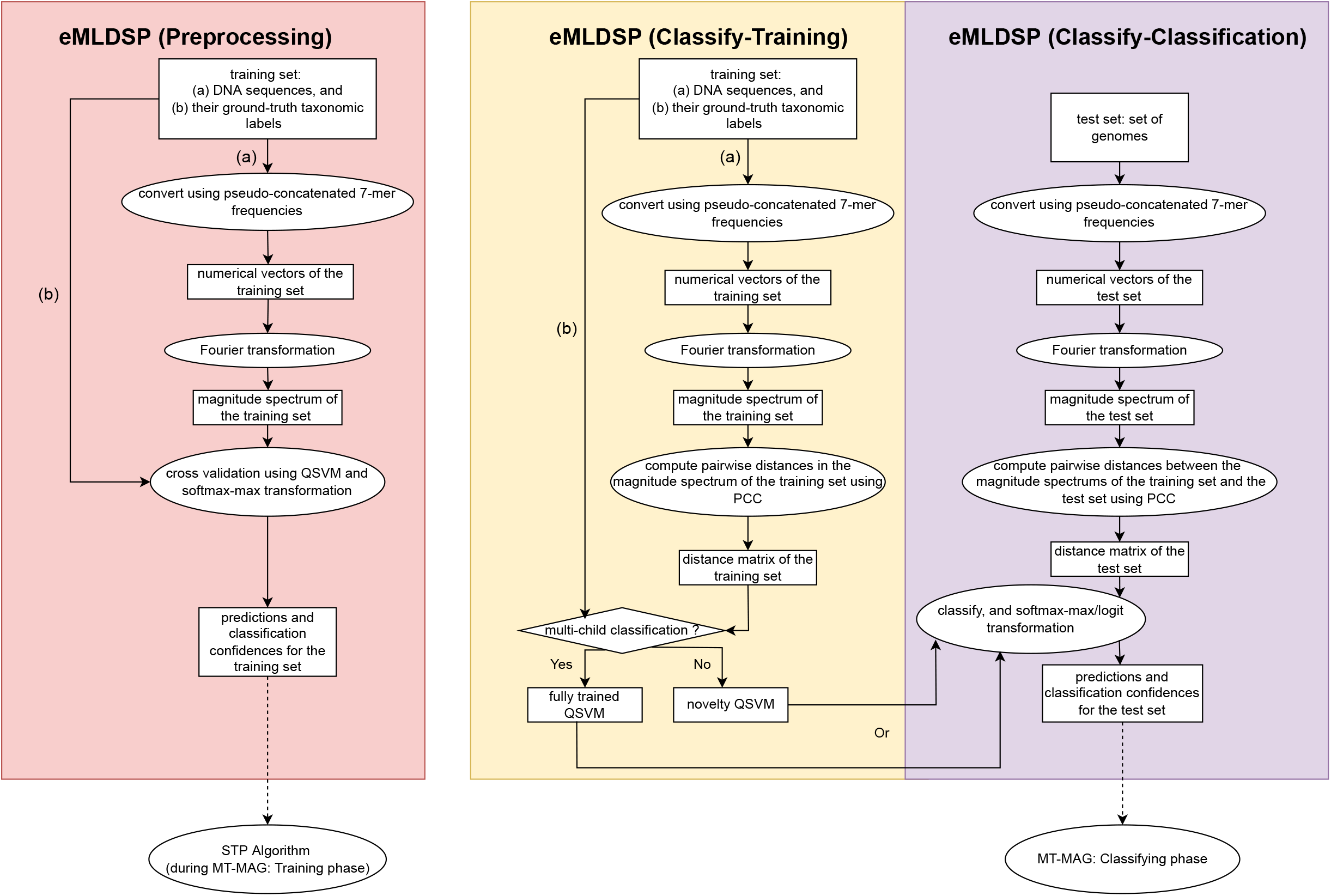
*Overview of eMLDSP*, including the main steps that comprise eMLDSP (Preprocessing) (pink box), eMLDSP (Classify-Training) (yellow box), and eMLDSP (Classify-Classification) (lavender box). Ellipses represent computation steps. Rectangles represent inputs to, and outputs from, computation steps. The diamond represents a condition checking. Note that the training dataset consists of (a) DNA sequences, together with (b) their taxonomic labels; and the inputs to eMLDSP (Preprocessing) and eMLDSP (Classify-Training) consists of the same training set. The output from eMLDSP (Preprocessing), consisting of predictions and classification confidences for the training set, is used for the STP algorithm in the MT-MAG Training phase (dotted arrow from the pink box), to calculate stopping thresholds. These stopping thresholds will then be used together with the output from eMLDSP (Classify-Classification) in the MT-MAG Classifying phase (dotted arrow from the lavender box), to determine final classification results.

The MLDSP implementation of the algorithms assumes that the input DNA sequences belong to multiple child taxa (*multi-child classification*). If this is the case, in the eMLDSP (Classify-Training) step, a Quadratic Support Vector Machine (QSVM) classifier called *fully trained QSVM* is trained, using the entire training set. In the eMLDSP (Classify-Classification) step, eMLDSP computes classifications (taxonomic assignments) for the DNA sequences in the test set by using the fully trained Quadratic Support Vector Machine (QSVM), and the classification confidences of these classifications using Platt scaling [31]. In contrast with its precursor, eMLDSP then applies five-fold cross-validation to obtain classifications, and uses a softmax-max transformation to compute classification confidences, for the entire training set.

Note that, when classifying a sequence belonging to a parent taxon, a single numerical classification confidence is computed for this classification, namely the confidence of classifying the sequence into the most likely child taxon of that parent taxon. This classification confidence is computed as the maximum of the posterior likelihoods over all child taxa. These results are later used for determining the stopping thresholds for each pair of parent and child taxon.

The case where the training/test sequences belong to a single child taxon (*single-child classification*) is not addressed by MLDSP. In this case, in the eMLDSP (Classify-Training) step, a QSVM classifier called *novelty QSVM* is trained, that uses the entire training set, and sets a fraction (default 10%) of the training set as a second child-class called “outlier taxon,” with the rest of the training set being referred to as the “original taxon.” In the eMLDSP (Classify-Classification) step, eMLDSP computes classifications for the DNA sequences in the test set by classifying using the novelty QSVM, and computes the classification confidences of these classifications by utilizing a normalizing logit transformation. The eMLDSP (Preprocessing) step is not applicable here, since there is no need for picking stopping thresholds in the case of single-child classifications.

#### The MT-MAG training phase and classifying phase

MT-MAG comprises two phases, training and classifying, as described below (see *S2 Appendix: Methods - MT-MAG algorithm* Section 3 and Section 4 for details of the two phases, and an additional optimization step that combines the two phases into a hybrid approach).

The **MT-MAG training phase** (of the training set comprising contigs in the case of Task 1 (sparse), respectively representative genomic fragments in the case of Task 2 (dense), together with their reference labels) comprises multiple training processes: For each parent taxon, after preparing the training set (discarding short sequences, handling imbalances in the dataset, etc.), two situations can occur, depending on the number of child taxa:

- *Multi-child classification*. In contrast to DeepMicrobes which uses a single stopping threshold, MT-MAG has multiple stopping thresholds, one for each parent-child pair. Concretely, MT-MAG determines a stopping threshold for every parent-child pair, based on the confidences calculated by eMLDSP (Preprocessing) with the training set as input. MT-MAG selects the stopping threshold from a list of candidate stopping thresholds, and searches for the stopping threshold *T* which results in the fewest number of contigs (resp. representative genomic fragments) with classification confidences lower than *T*, while at the same time resulting in the classification accuracies of the other contigs (resp. representative genomic fragments) being higher than the value of a user-specified accuracy parameter. More specifically, a stopping threshold is the result of subtracting a “variability” parameter from the maximum of *(a)* the minimum of the candidate thresholds (numbers between 0 and 1) that result in a “constrained accuracy” being greater than the value of a user-specified parameter (default: 90%), and *(b)* the average of classification confidences for the contigs (resp. representative genomic fragments) with correct eMLDSP (Preprocessing) classifications. Subsequently, a QSVM classifier (the fully trained QSVM) is trained with the entire training set of this parent taxon, as part of the eMLDSP (Classify-Training) step.
- *Single-child classification*. A QSVM classifier called *novelty QSVM* is trained in the eMLDSP (Classify-Training) step. The novelty QSVM sets a fraction of the contigs (resp. representative genomic fragments) in the training set as a second child-class, called *outlier taxon*. The default fraction is set to 10%.

The **MT-MAG classifying phase** (of the test set comprising test genomes with known reference labels, or unknown genomes) proceeds as follows. When, in the process of hierarchically-structured local classification, MT-MAG has classified a test/unknown genome into a parent taxon, and attempts to classify it further into one of its child taxa, two possibilities can occur:

- *Multi-child classification*. If the parent taxon has multiple child taxa, then the fully trained QSVM is used to classify the test/unknown genome into one of the child taxa, and this result is also used to compute a classification confidence as part of the eMLDSP (Classify-Classification) step. If this classification confidence is below the stopping threshold for this parent-child pair, then this classification is considered uncertain, and no further attempts are made to classify this test/unknown genome from the child taxon into its own child taxa.
- *Single-child classification*. If the parent taxon has a single child taxon, then the novelty QSVM is used to classify the test/unknown genome into either the child taxon or the outlier taxon as part of the eMLDSP (Classify-Classification) step, and the result is used to compute a classification confidence. If the output is the outlier taxon, then this classification is considered uncertain and no further classifications are attempted.

Given a test/unknown genome, the output of MT-MAG is either *(i)* a complete classification path down to the Species level, if all the intermediate classification confidences are greater than or equal to the stopping thresholds, or *(ii)* a partial classification path, down to the lowest taxonomic rank with a high enough classification confidence. In either case, the output of MT-MAG also includes the classification confidence for each taxon along the classification path.

## MT-MAG multi-child classification pipeline

Fig 3 illustrates the MT-MAG training phase and classifying phase for classifying two genomes belonging to a given parent taxon, into one of its two child taxa (multi-child classification). In the MT-MAG training phase, the classifications and classification confidences outputted in eMLDSP (Preprocessing) for the training set from all folds are used for determining the stopping thresholds for every child taxon of this parent taxon. Furthermore, in eMLDSP (Classify-Training), a fully trained QSVM is trained by using the entire training data. In the MT-MAG classifying phase, the test set is given as the input to eMLDSP (Classify-Classification), together with the fully trained QSVM from the training phase. For each genome in the test set, the output of eMLDSP (Classify-Classification) is a classification, together with a numerical confidence (between 0 and 1) of that classification. This classification confidence is then compared with the stopping threshold of that parent taxon and child taxon pair. If the classification confidence is lower than the stopping threshold, then the output is “uncertain classification,” and further classification into children of this child taxon will not be attempted. See *S2 Appendix: Methods - MT-MAG algorithm* Section 2 for a visual illustration of the single-child classification scenario, similar to Fig 3.

**Fig 3.**
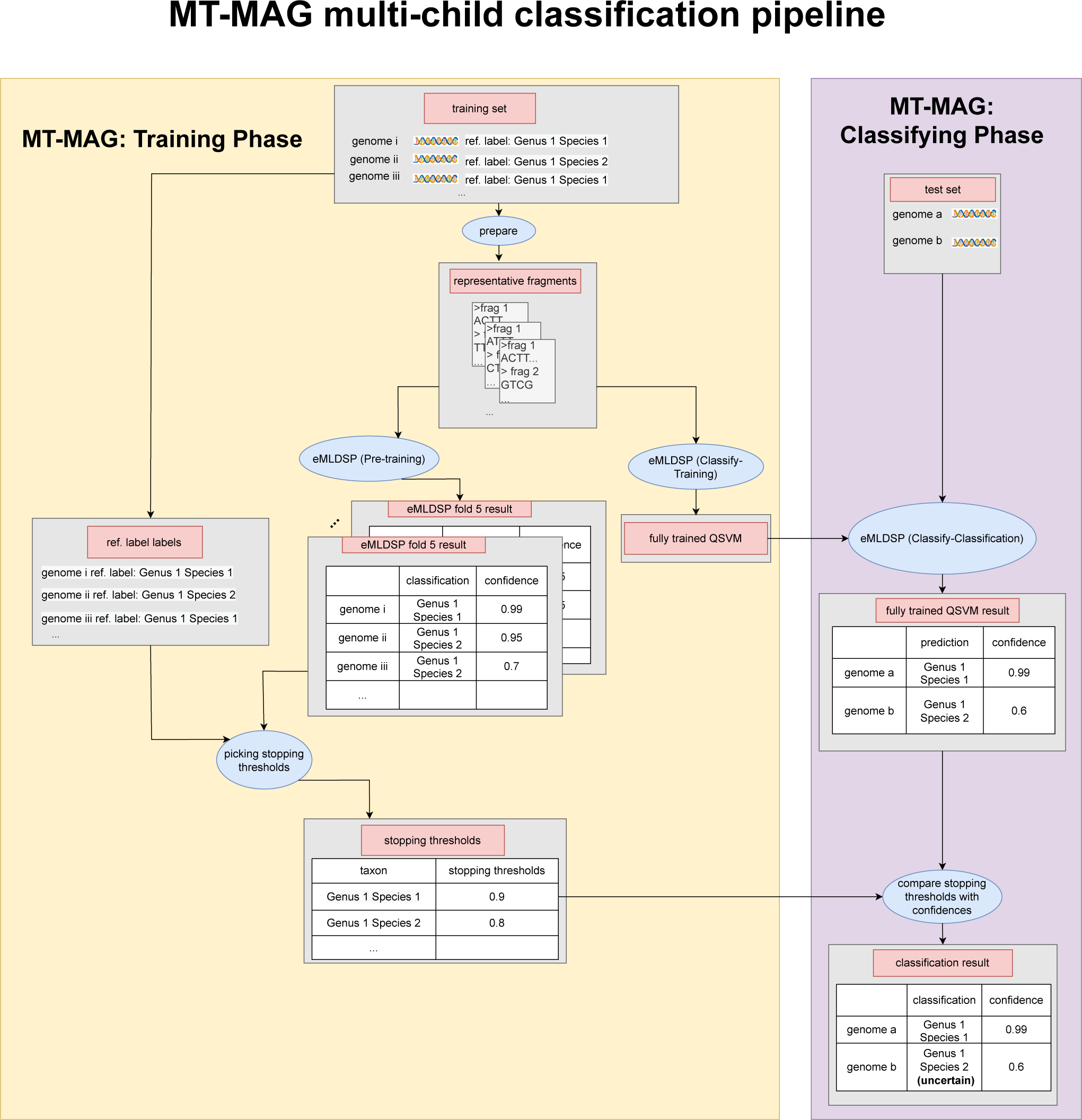
*MT-MAG pipeline for classifying two genomes, genome a and genome b, from the parent taxon Genus* 1 *into its two child taxa, Species* 1, *and Species* 2 *(multi-child classification)*. Blue ellipses represent computation steps. Gray rectangles represent inputs to, and outputs from, computation steps. In the MT-MAG training phase (yellow box), the training set is prepared and given as the input to eMLDSP (Preprocessing).

### Performance metrics

In this section, we define the terminology and the performance metrics used to discuss and assess the performance of MT-MAG’s classification of test genomes.

A classification of a genome *x* from taxonomic rank *tr*_1_ to taxonomic rank *tr*_2_, is called *a classification at tr*_2_. Given a taxonomic rank *tr*, we call the classification of *x* a *complete classification at tr* if, for each classification on the classification path for *x*:

- *if multi-child*, the classification confidence of classifying *x* at any taxonomic level higher than and including *tr*, is greater than or equal to the respective stopping threshold, and
- *if single-child*, it is classified as the original single-child taxon.

The classification of *x* is called an *uncertain classification at tr* if:

a. for the classification at *tr*,
  – *if multi-child*, the confidence of the classification at *tr* is strictly less than the stopping threshold of this parent-child pair,
  – *if single-child*, it is classified to the outlier taxon;
b. for each classification at ranks higher than *tr*,
  – *if multi-child*, the confidence of the classification is greater than or equal to its corresponding stopping threshold,
  – *if single-child*, it is classified to the original single-child taxon.

There are three possible causes for the *uncertain classification* of *x* to *tr*. First, *x* belongs to *tr*, but MT-MAG fails to confidently classify it to *tr*. Second, *x* belongs to another existing taxon with the same parent taxon as *tr*, and MT-MAG successfully identifies this. Third, *x* belongs to another taxon with the same parent taxon as *tr*, that does not exist in the training set. Here, MT-MAG’s uncertain classification suggests the discovery of a new taxon with the same parent taxon as *tr*.

If the classification is uncertain a rank higher than *tr*, we call it an *unattempted classification at tr*. At the end of the MT-MAG classifying phase for an input genome *x*, if the classification of *x* is uncertain at any taxonomic rank lower than the first non-root rank, then we say that *x* is *partially classified*. On the other hand, if the output of the classifying phase is that *x* is completely classified at the lowest taxonomic rank (herein Species), then we say that *x* is *completely classified*.

We note that for a given test genome, the output of its classification at rank *tr* can be only one of the following: complete classification at *tr* (if classifications all the way down to *tr* exceed their thresholds), uncertain classification at *tr* (if classifications at all higher ranks exceed their thresholds, but the classification at *tr* is below the threshold), or unattempted classification at *tr* (if the classification at any rank higher than the one right above *tr* is below the threshold).

We also say that a classification of *x* is a *correct classification down to tr* if it is a complete classification at *tr*, and a correct classification at all taxonomic ranks higher than, and including, *tr*. By definition, all genomes have correct classifications down to the root.

Lastly, the *length of the classification path of x* is defined as either *(i)* the number of taxa in the taxonomy, if *x* is completely classified at the lowest taxonomic rank, or *(ii)* the number of taxa predicted before the uncertain classification, if the output is “uncertain classification.”

As an example, in Fig 4, the classification of genome *x* is a complete classification at rank 1, an uncertain classification at rank 2, and an unattempted classification at rank 3. In addition, if the reference taxon of genome *x* at rank 1 is “rank 1 group 1”, then the classification of genome *x* is a correct classification at rank 1, as well as a correct classification down to rank 1. Note that, for each classification of a parent taxon, the number of stopping thresholds equals the number of that parent’s child taxa. In contrast, each such classification has associated with it a single classification confidence, that of classifying the genome into a single, “best-guess,” child taxon.

**Fig 4.**
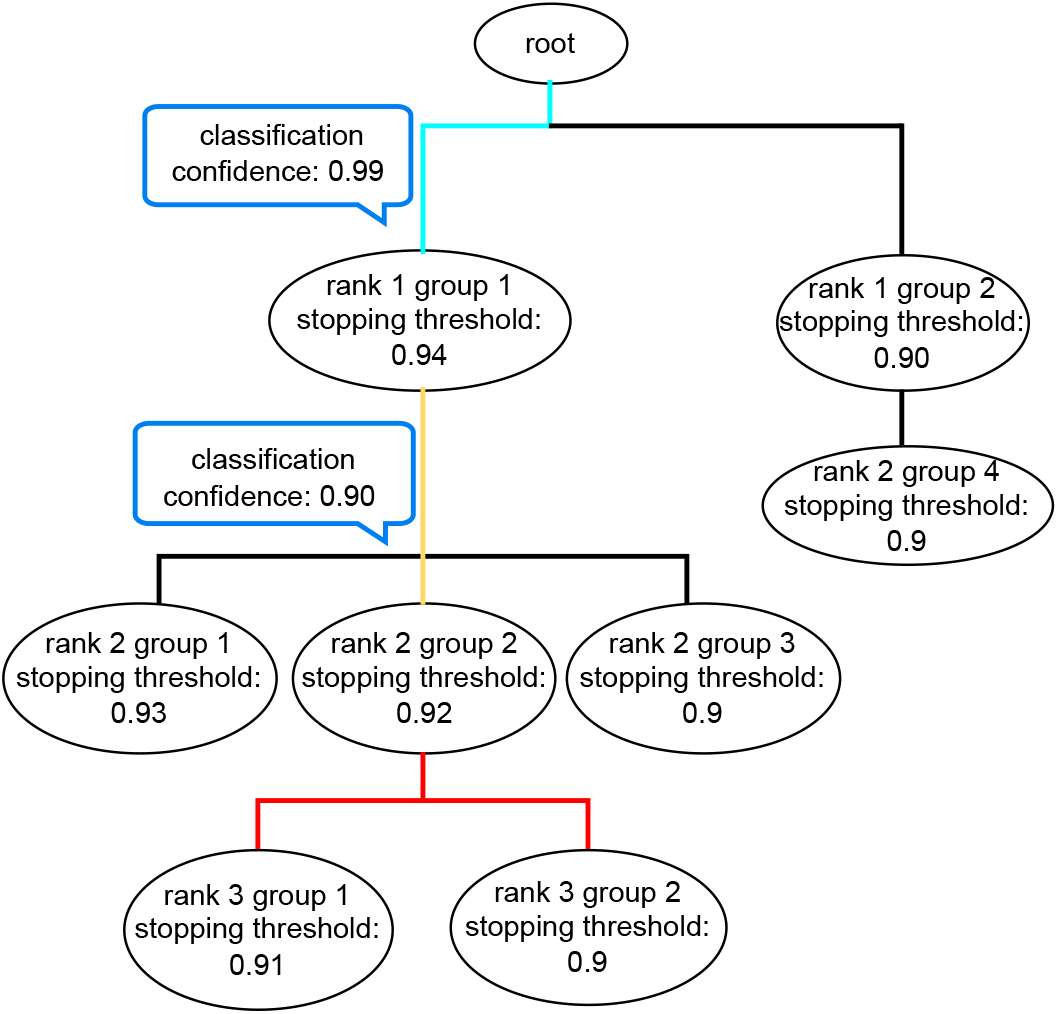
Example of the classification path for a genome *x*. The pre-calculated stopping thresholds are listed under the corresponding taxon labels. The classification confidences are listed inside blue-bordered rectangles. MT-MAG classifies *x* from root into “rank 1 group 1” with confidence 0.99, which is greater than the stopping threshold for “rank 1 group 1” (0.94), so MT-MAG continues its classification for *x*. In the next iteration MT-MAG classifies *x* from “rank 1 group 1” into “rank 2 group 2” with confidence 0.90, but since this is below the stopping threshold of the parent into its child “rank 2 group 2” (0.92), this classification is deemed “uncertain” and MT-MAG does not attempt further classifications. The path in cyan indicates complete classification(s), the path in yellow indicates uncertain classification(s), and the part in red indicates unattempted classifications.

With this terminology, for a given taxonomic rank *tr*_*j*_ in a list of increasingly lower taxonomic ranks *tr*_0_, *tr*_1_, …, *tr*_*i*_, where *tr*_0_ is the root, we define the following performance metrics (the subscript *g* indicates that these metrics refer to genomes):

- *CA*_*g*_(*tr*_*j*_) (constrained accuracy): the proportion of the test genomes with correct classifications down to *tr*_*j*_, to the test genomes with complete classifications at *tr*_*j*_.
- *AA*_*g*_(*tr*_*j*_) (absolute accuracy): the proportion of the test genomes with correct classifications down to *tr*_*j*_, to all test genomes.
- *WA*_*g*_(*tr*_*j*_) (weighted classification accuracy) is the ratio between the weighted sum computed as described below, and the number of test genomes. Given a test genome with a classification path of length 0 ≤ *h* ≤ *i*, (where *i* is the maximum length of a classification path), its weight is:
  – If it has a correct classification down to *tr*_*j*_ (i.e., complete classification at *tr*_*j*_, *and* it a correct classification at *tr*_*j*_), then the weight is 1.
  – If, on the other hand, there exists a 0 *< p < j*, such that the test genome has a correct classification down to *tr*_*p*_, but does not have a correct classification at *tr*_*p*+1_ (the latter condition avoids double counting), then the weight of the test genome is

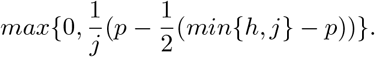

With the formula above, genomes that do not have correct classifications down to any taxonomic rank below the root will have weight 0 (the underlying assumption is that the test genomes are always assumed to belong to the root). The term *p* is intended to give positive weights to the correct classifications down to *tr*_*p*_. The term 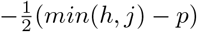 is intended to penalize incorrect classifications after taxonomic rank *tr*_*p*_ by 0.5, while taking the maximum between the calculated result and 0 is meant to disallow negative weights.

This weighting scheme reflects the fact that partial classifications at different ranks are not equally informative. For example, a correct classification of a test genome down to the Phylum level is less informative than a correct classification of a test genome down to the Genus level.

Hereafter, *weighted classification accuracy* will sometimes be called just *weighted accuracy*.

- *CR*_*g*_(*tr*_*j*_) (complete classification rate): the proportion of the test genomes with complete classifications at *tr*_*j*_, to all test genomes.

Given a taxonomic rank *tr*_*j*_, the three accuracies *CA*_*g*_(*tr*_*j*_), *AA*_*g*_(*tr*_*j*_) and *WA*_*g*_(*tr*_*j*_) are numbers between 0 and 1, with *CA*_*g*_(*tr*_*j*_) ≥ *AA*_*g*_(*tr*_*j*_), and where higher values indicate better performance. The complete classification rate *CR*_*g*_(*tr*_*j*_) is a number between 0 and 1, and a higher value indicates a higher proportion of genomes that are completely classified at *tr*_*j*_. See *S3 Appendix: Results* Section 3 for the formal definitions of these performance metrics.

## Results

In this section, we first present a detailed analysis of the features of MT-MAG: *(i)* the capability to classify a DNA sequence at all taxonomic ranks, *(ii)* the capability to output an interpretable classification confidence for the classification at each taxonomic rank along the classification path, and *(iii)* the capability to output a “partial classification” path when the classification confidence of a classification does not meet a given threshold. Second, since DeepMicrobes is able to classify only at the Species level, we summarize the results of a comparative analysis of the performance of MT-MAG with that of DeepMicrobes at the Species level. Note that, in the comparative analysis, some genomes were excluded from the test sets as follows: the genomes for which the GTDB taxon is “unnamed species” were excluded in both Task 1 and Task 2, and the genomes for which the reference GTDB species did not exist in the training set were excluded in Task 1 (see *S3 Appendix: Results* Section 2).

Experiments for DeepMicrobes were run using Vector Institute’s Vaughan cluster with GPUs of rtx6000, t4v1,p100 and t4v2 with 24 GB memory. Experiments for MT-MAG were run on a workstation with x86_64 CPU.

## MT-MAG features

### Classifications at all taxonomic ranks

In contrast with DeepMicrobes which only classifies reads at the Species level, a significant feature of MT-MAG is its capability to classify genomes at all taxonomic ranks. Table 2 provides a summary of MT-MAG’s performance metrics, at all taxonomic ranks, for both Task 1 (sparse) and Task 2 (dense). Table 3 provides a summary of the percentages of the test sequences completely classified by MT-MAG vs. classified by DeepMicrobes, at all taxonomic ranks.

**Table 2.** Summary of MT-MAG performance metrics at all taxonomic ranks, for Task 1 (sparse) and Task 2 (dense): constrained accuracy *CA*_*g*_(*tr*), absolute accuracy *AA*_*g*_(*tr*), weighted accuracy *WA*_*g*_(*tr*), and complete classification rate *CR*_*g*_(*tr*) (higher is better).

| Task ID | Taxonomic Rank | $CA_g(tr)(\%)$ | $AA_g(tr)(\%)$ | $WA_g(tr)(\%)$ | $CR_g(tr)(\%)$ |
| --- | --- | --- | --- | --- | --- |
| Task 1 (sparse) | Phylum | 100.00 | 93.80 | 93.80 | 93.80 |
|  | Class | 100.00 | 93.80 | 93.80 | 93.80 |
|  | Order | 99.56 | 91.99 | 93.13 | 92.40 |
|  | Family | 100.00 | 82.67 | 90.52 | 82.67 |
|  | Genus | 99.39 | 71.53 | 86.68 | 71.96 |
|  | Species | 81.53 | 57.60 | 80.75 | 70.65 |
|  | **Average** | 96.75 | 81.90 | 89.78 | 84.21 |
| Task 2 (dense) | Domain | 99.66 | 96.13 | 95.97 | 96.46 |
|  | Phylum | 97.50 | 83.82 | 88.34 | 83.87 |
|  | Class | 97.74 | 81.10 | 85.62 | 82.98 |
|  | Order | 97.96 | 79.56 | 83.90 | 81.22 |
|  | Family | 98.06 | 78.12 | 82.59 | 79.67 |
|  | Genus | 96.70 | 67.96 | 79.95 | 70.28 |
|  | Species | 98.63 | 63.87 | 77.59 | 64.75 |
|  | **Average** | 98.03 | 78.36 | 84.85 | 79.89 |

**Table 3.** Summary of proportion of test sequences completely classified by MT-MAG (quantified as *CR*_*g*_(*tr*)) vs. classified by DeepMicrobes (quantified as *CR*_*r*_), at all taxonomic ranks. A higher *CR*_*g*_(*tr*) (respectively *CR*_*r*_) is better, as it signifies that a higher proportion of genomes (resp. reads) have been completely classified (resp. classified). Dash denotes not applicable.

| Task ID | Taxonomic Rank | MT-MAG $CR_g(tr)(\%)$ | DeepMicrobes $CR_r(\%)$ |
| --- | --- | --- | --- |
| Task 1 (sparse) | Phylum | 93.80 | — |
|  | Class | 93.80 | — |
|  | Order | 92.40 | — |
|  | Family | 82.67 | — |
|  | Genus | 71.96 | — |
|  | Species | 70.65 | 45.02 |
| Task 2 (dense) | Domain | 96.46 | — |
|  | Phylum | 83.87 | — |
|  | Class | 82.98 | — |
|  | Order | 81.22 | — |
|  | Family | 79.67 | — |
|  | Genus | 70.28 | — |
|  | Species | 64.75 | 49.88 |

In Task 1 (sparse) MT-MAG accuracies range between 82% and 97% (see Table 2).

Specifically, the MT-MAG constrained accuracies *CA*_*g*_(*tr*) are above 99% at all taxonomic ranks, except at the Species level *CA*_*g*_(*Species*), where they drop to 81.53%. The increase in the number of incorrect classifications at the Species level explains, in part, the 5.93% drop in weighted accuracy *WA*_*g*_(*tr*) from the Genus to the Species level. In addition, due to its partial classification capability, MT-MAG is able to completely classify 93.80% of the test genomes to the Phylum and Class levels, 92.40% to the Order level, 82.67% to the Family level, and 71.96% to the Genus level, with 70.65% of the test genomes being completely classified to the Species level (see Table 3). In contrast, in Task 1, DeepMicrobes classifies only 45.02% of the test reads at the Species level, and it does not assess other taxonomic levels.

In Task 2 (dense) MT-MAG has a satisfactory performance all around, with constrained accuracies *CA*_*g*_(*tr*) above 96% at all taxonomic ranks (see Table 2). In addition, due to its partial classification capability, MT-MAG completely classifies 83.87% of the test genomes to the Phylum level, 82.98% to the Class level, 81.22% to the Order level, 79.67% to the Family level, and 70.29% to the Genus level, with 64.75% of the test genomes being completely classified to the Species level (see Table 3). In contrast, in Task 2, DeepMicrobes only classifies 49.88% of the test reads to the Species level, and does not assess other taxonomic levels.

Overall, for the two benchmarking datasets, MT-MAG completely classifies an average of 67.70% of the test sequences (to the Species level). In addition, MT-MAG provides partial classifications for the majority of the remaining sequences. This results in 93.80% of genomes analyzed in Task 1 (sparse) and 96.46% of genomes analyzed in Task 2 (dense) being partially classified or completely classified. In particular, due to its partial classification capability, MT-MAG completely classifies on average 88.84% of the test sequences to the Phylum level, 88.39% to the Class level, 86.81% to the Order level, 81.17% to the Family level, and 71.13% to the Genus level. Moreover, MT-MAG provides additional information for sequences that it could not classify at the Species level, resulting in an average of the partial or complete classification of 95.13% (i.e., (93.80%+96.46%)/2=95.13%), of the genomes in the datasets analyzed.

### Numerical classification confidences

In addition to the final classification path, MT-MAG also outputs numerical classification confidences along the classification path, indicating how confident MT-MAG is in the classification, at each taxonomic rank. For example, the final classification path for genome *x*, illustrated in Fig 4 is interpreted as MT-MAG being 99% confident in classifying *x* from “root” to “rank 1 group 1,” and 90% confident in classifying *x* from “rank 1 group 1” to “rank 2 group 2.” However, since the confidence of the latter classification is strictly less than the pre-calculated stopping threshold of 92%, this classification is deemed “uncertain” and no further classifications are attempted for genome *x*.

As an example of a complete classification down to the Species level, in Task 2 (dense) the final classification path for genome hRUG888 is “Domain Bacteria (confidence 97%) → Phylum Bacteroidota (confidence 97%) → Class Bacteroidia (confidence 100%) → Order Bacteroidales (confidence 100%) → Family Muribaculaceae (confidence 99%) → Genus *Sodaliphilus* (confidence 99%) → Species *Sodaliphilus* sp900314215 (confidence 99%).” As an example of a partial classification path, the final classification path for genome RUG412 is “Domain Bacteria (confidence 93%) → Phylum Bacteroidota (confidence 100%) → Class Bacteroidia (confidence 100%) → Order Bacteroidales (confidence 100%) → Family Muribaculaceae (confidence 98%) → Genus *Sodaliphilus* (confidence 99%) → Species *Sodaliphilus* sp900318645 (uncertain).” The last output means that MT-MAG is uncertain regarding its classification of RUG412 from Genus *Sodaliphilus* into Species *Sodaliphilus* sp900318645.

### Species level comparison of MT-MAG with DeepMicrobes

In this section, we compare the performance of MT-MAG against the performance of DeepMicrobes at the Species level, the only taxonomic rank at which DeepMicrobes classifies. The performance metrics we define here are used to assess the quality of DeepMicrobes’s classification, and are defined analogously to the performance metrics for MT-MAG. The subscript *r* indicates that these metrics refer to reads, and the exact definitions of the terms used can be found in, “Materials: Datasets and task description”. These performance metrics are:

- *CA*_*r*_ (constrained accuracy): the proportion of correctly classified test reads, to classified test reads.
- *AA*_*r*_ (absolute accuracy): the proportion of correctly classified reads, to all test reads.
- *WA*_*r*_ (weighted classification accuracy): the proportion of correctly classified reads. Note that in this case *WA*_*r*_ coincides with *AA*_*r*_, since DeepMicrobes does not provide any classification at ranks other than Species. (hereafter, *weighted classification accuracy* will sometimes simply be called *weighted accuracy*.)
- *CR*_*r*_ (classified rate): the proportion of classified test reads to all test reads. Note the difference in the definition between *CR*_*g*_(*tr*) for MT-MAG (complete classification rate for genomes, at rank *tr*), and *CR*_*r*_ (classified rate for reads, at the Species level) for DeepMicrobes.

The three accuracies *CA*_*r*_, *AA*_*r*_ and *WA*_*r*_ are numbers between 0 and 1, with *CA*_*r*_ ≥ *AA*_*r*_, and where higher values indicate better performance. The classified rate *CR*_*r*_ is a number between 0 and 1, and a higher value indicates a higher proportion of classified reads, at the Species level (for exact definitions, see *S3 Appendix: Results* Section 3).

Since DeepMicrobes only makes classifications at the Species level, to compare its performance with that of MT-MAG, we set the parameter *tr* (taxonomic rank) to Species in MT-MAG, and proceeded to compare *CA*_*r*_ with *CA*_*g*_(Species), *AA*_*r*_ with *AA*_*g*_(Species), *WA*_*r*_ with *WA*_*g*_(Species), and *CR*_*r*_ with *CR*_*g*_(Species).

Of all the metrics we defined, we posit that the most informative metric for comparing MT-MAG with DeepMicrobes is the *weighted (classification) accuracy* at the Species level. Indeed, in the case of MT-MAG, *WA*_*g*_(Species) combines, into a single numerical indicator, the information on the proportion of genomes that MT-MAG correctly classifies together with that of genomes that it partially classifies. In the case of DeepMicrobes, *WA*_*r*_ combines the information on the proportion of reads that it correctly classifies together with that of reads that it is unable to classify. In addition to this main comparison performance metric, and for a more nuanced discussion, in the following we also compare the other performance metrics, namely *CA*_*r*_ with *CA*_*g*_(Species), *AA*_*r*_ with *AA*_*g*_(Species), and *CR*_*r*_ with *CR*_*g*_(Species).

Table 4 summarizes the MT-MAG and DeepMicrobes constrained accuracies, absolute accuracies, and *weighted accuracies*, as well as the complete classification rates of MT-MAG, respectively the classified rates of DeepMicrobes.

**Table 4.** Summary of MT-MAG and DeepMicrobes accuracy statistics, as well as the complete classification rates of MT-MAG and the classified rates of DeepMicrobes. The inputs are genomes in the case of MT-MAG, and reads in the case of DeepMicrobes. A higher value indicates better performance (in boldface). The metric that best captures the performance of the methods is the weighted accuracy (in blue), since this metric combines information about sequences that have been completely classified with information about the sequences that have not been completely classified to the Species level.

| Task ID | Metric | MT-MAG(%) | DeepMicrobes(%) |
| --- | --- | --- | --- |
| Task 1 (sparse) | $CA_g(\text{Species})/CA_r$ | 81.53 | **93.14** |
| | $AA_g(\text{Species})/AA_r$ | **57.60** | 41.94 |
| | $WA_g(\text{Species})/WA_r$ | **80.75** | **41.94** |
| | $CR_g(\text{Species})/CR_r$ | **70.65** | 45.02 |
| Task 2 (dense) | $CA_g(\text{Species})/CA_r$ | **98.63** | 93.87 |
| | $AA_g(\text{Species})/AA_r$ | **63.87** | 46.82 |
| | $WA_g(\text{Species})/WA_r$ | **77.59** | **46.82** |
| | $CR_g(\text{Species})/CR_r$ | **64.75** | 49.88 |

For Task 1 (sparse), as seen in Table 4, MT-MAG demonstrates significantly better overall performance than DeepMicrobes, with the weighted accuracy of MT-MAG being 38.81% higher than that of DeepMicrobes. Regarding other performance metrics, the constrained accuracy of DeepMicrobes 11.61% higher than that of MT-MAG, the absolute accuracy for MT-MAG is 15.66% higher than that of DeepMicrobes, and the complete classification rate of MT-MAG is 25.63% higher than the classified rate of DeepMicrobes. The latter indicates that MT-MAG completely classifies significantly more sequences than DeepMicrobes, though DeepMicrobes demonstrates a slightly higher constrained classification accuracy for the classified sequences.

For Task 2 (dense), as seen in Table 4, MT-MAG demonstrates significantly better overall performance than DeepMicrobes, with the weighted accuracy of MT-MAG being 30.77% higher than that of DeepMicrobes. Comparing the other performance metrics, the constrained accuracy of MT-MAG is 4.76% higher than that of DeepMicrobes, the absolute accuracy for MT-MAG is 17.05% higher than that of DeepMicrobes, and the complete classification rate of MT-MAG is 14.87% higher than the classified rate of DeepMicrobes. This indicates that MT-MAG not only completely classifies significantly more sequences than DeepMicrobes, but also demonstrates a slightly higher MT-MAG classification accuracy for the completely classified sequences.

Overall, for Task 1 (sparse) and Task 2 (dense), MT-MAG outperforms DeepMicrobes by an average of 34.79% in weighted accuracy. In addition, MT-MAG is able to completely classify an average of 67.70% of the sequences at the Species level, the only comparable taxonomic rank of DeepMicrobes, which only classifies 47.45%.

## Discussion

We proposed MT-MAG, a novel *alignment-free* and *genetic marker-free* software tool that uses machine learning to obtain taxonomic assignments of metagenome-assembled genomes. MT-MAG employs *k*-mer frequencies (here in *k* = 7) as the only feature used to distinguish between DNA sequences. MT-MAG has a partial classification option, whereby MT-MAG outputs a partial classification at a higher taxonomic rank for the MAGs that it cannot confidently classify to the lowest taxonomic rank (e.g., Species). In addition, MT-MAG outputs interpretable numerical classification confidences of its classifications, at each taxonomic rank.

To assess the performance of MT-MAG, we defined a “weighted accuracy,” with a weighting scheme reflecting the fact that partial classifications at different ranks are not equally informative. Compared with DeepMicrobes (the only other machine learning tool for taxonomic assignment of metagenomic data, with confidence scores), for the two datasets analyzed (genomes from human gut microbiome species, respectively bacterial and archaeal genomes assembled from cow rumen metagenomic sequences), MT-MAG outperforms DeepMicrobes by an average of 34.79% in weighted accuracy. In addition, MT-MAG is able to completely classify an average of 67.70% of the sequences at the Species level, the only comparable taxonomic rank of DeepMicrobes, which only classifies 47.45%. Moreover, a significant feature of MT-MAG is that it provides additional information for the sequences that are not completely classified at the Species level. This results in 95.13% of the genomes analyzed being either partially classified or completely classified, averaged over the two datasets analyzed. In particular, due to its partial classification capability, MT-MAG completely classifies, on average, 88.84% of the test genomes to the Phylum level, 88.39% to the Class level, 86.81% to the Order level, 81.17% to the Family level, and 71.13% to the Genus level.

In addition, MT-MAG is able to be run using a reasonable amount of random access memory (RAM) and on-chip memory. Indeed, for the computational experiments in this study, MT-MAG was run on a workstation where each CPU had only 0.02 GB of on-chip RAM. DeepMicrobes could not be run on this workstation, as it required significantly more memory resources (several GPUs, each with 24 GB of on-chip memory).

Limitations of MT-MAG include the fact that, being a supervised machine learning classification algorithm, its performance relies on the availability of reference taxonomic labels for the DNA sequences in the training set. In addition, any incorrect or unstable reference labels in the training set may cause erroneous future classifications. This limitation could be addressed, e.g., by extending the supervised machine learning approach to semi-supervised machine learning (where some, but not all, information about the training set is available), or even to unsupervised machine learning (where the training process does not require any reference taxonomic labels, see, e.g., [32]).

Second, even though MT-MAG significantly outperforms DeepMicrobes in Task 1 (sparse training set) and Task 2 (dense training set) in weighted accuracy, there is still room for improvement in accuracies and complete classification rates. An analysis of the Task 1 (sparse) training set suggests two possible reasons contributing to incorrect classifications. One reason is the fact that the training set was the HGR database, which constitutes a very small subset of the GTDB taxonomy, in terms of both the number of representative genomes and of coverage of the GTDB taxonomy. This could be addressed by requiring a specific level of coverage for known taxa, to ensure that feature characteristics are reasonably well-represented. Another reason is the fact that, due to computational requirements of MLDSP, the training set had to exclude any contigs shorter than 5,000 bp, and this selection process resulted in the removal of 93% of the available basepairs. This could be addressed by finding ways to relax the selection criteria for the training set, to allow more sequences to participate in the training process without compromising the classification performance.

Third, the interpretability of classification could be further enhanced by exploring the last layer of the classifier. For example, the process of computing classification confidences could be used to identify pairs of child taxa that are difficult to distinguish from each other, which could potentially be biologically relevant. In addition, while single-child cases are few in the case of real DNA datasets, we note that their classification confidences are computed via a transformation of the distances between a test sequence and decision boundaries in the feature space into a valid probability distribution. To enhance the interpretability of these single-child class confidences, one could consider applying more interpretable training process and transformations such as those proposed in [33, 34].

Fourth, the classification accuracy and computational efficiency of MT-MAG could be further improved by taking advantage of user-provided information, so that the computation does not always start from the root of the taxonomy. For example, if the user already knows that an input genome belongs to Class Bacteroidia, then MT-MAG could bypass the higher taxonomic ranks and start its training and classifying phases at the Class-to-Order level directly.

Fifth, when defining the weighted accuracy for a classification at given taxonomic rank *tr* (i.e., *WA*_*g*_(*tr*_*i*_)), the weights used in this computation can be further refined, to reflect the dataset analyzed. Recall that the intent of defining a weighted classification accuracy was to account for the fact that partial classifications of a genome at different ranks are not equally informative. For example, a partial classification of a genome down to the Genus level is intuitively more informative than a partial classification to, say, the Phylum level, and this is quantified as follows in the definition of weighted accuracy. The root is assigned weight 0, the last taxonomic rank with a correct classification is assigned weight 1, and intermediate taxonomic ranks are assigned weights that increase in *equal* fractional increments, from the root to the last correctly classified rank. However, this assumption of equal increments at each intermediate rank could be inadequate if, e.g., some of the intermediate taxonomic ranks are missing from the path. In such cases, the individual weights of taxonomic ranks could be defined as being different, with each weight corresponding to the amount of information that a classification at that rank contributes.

Lastly, even though MT-MAG achieves superior performance on the datasets analyzed in this paper, it would be desirable to obtain mathematical proofs of the optimality of the classifier, such as the Bayes optimality proofs in [35].

## Acknowledgments

We thank DeepMicrobes’s co-author Qiaoxing Liang for the assistance with DeepMicrobes, Dr. Gurjit Randhawa for the guidance on MLDSP, Fatemeh Alipour and Pablo Millan Arias for replicating the results and proofreading the manuscript. Resources used in preparing this research were provided, in part, by the Province of Ontario, the Government of Canada through CIFAR, and the Vector Institute.

## Tools and data availability

MT-MAG presented in this paper and scripts for Task 1 (simulated/sparse) and Task 2 (real/dense) are fully available at https://github.com/wxli0/MT-MAG.git. The subroutine eMLDSP is available at https://github.com/wxli0/MLDSP.git. The scripts for running DeepMicrobes for benchmarking are available at https://github.com/wxli0/DeepMicrobes.git.

## Supporting information

*S1 Appendix: Materials - Datasets and task description*

*S2 Appendix: Methods - MT-MAG algorithm*

*S3 Appendix: Results*

## Notes

### Competing Interest Statement

The authors have declared no competing interest.

### Summary of Updates

Main manuscript and supplemental files updated.

## References

1. Sharon I, Banfield JF. Genomes from metagenomics. Science. 2013;342(6162):1057–8. doi:10.1126/science.1247023.

2. Parks DH, Rinke C, Chuvochina M, Chaumeil PA, Woodcroft BJ, Evans PN, Hugenholtz P, Tyson GW. Recovery of nearly 8,000 metagenome-assembled genomes substantially expands the tree of life. Nature Microbiology. 2017;2(11):1533–42. doi:10.1038/s41564-017-0012-7.

3. Murali A, Bhargava A, Wright ES. IDTAXA: A novel approach for accurate taxonomic assignment of microbiome sequences. Microbiome. 2018;6(1):1–4. doi:10.1186/s40168-018-0521-5.

4. Frioux C, Singh D, Korcsmaros T, Hildebrand F. From bag-of-genes to bag-of-genomes: metabolic modelling of communities in the era of metagenome-assembled genomes. Computational and Structural Biotechnology Journal. 2020;18:1722–34. doi:10.1016/j.csbj.2020.06.028.

5. Parks DH, Chuvochina M, Waite DW, Rinke C, Skarshewski A, Chaumeil PA, Hugenholtz P. A standardized bacterial taxonomy based on genome phylogeny substantially revises the tree of life. Nature Biotechnology. 2018;36(10):996–1004. doi:10.1038/nbt.4229.

6. Parks DH, Chuvochina M, Chaumeil PA, Rinke C, Mussig AJ, Hugenholtz P. A complete Domain-to-Species taxonomy for Bacteria and Archaea. Nature Biotechnology. 2020;38(9):1079–86. doi:10.1038/s41587-020-0501-8.

7. Derrick E Wood, Jennifer Lu and Ben Langmead Improved metagenomic analysis with Kraken 2 Genome Biology. 2019;20(1):1–13.

8. Mock F, Kretschmer F, Kriese A, Böcker S, Marz M. BERTax: taxonomic assignment of DNA sequences with deep neural networks. BioRxiv. 2021. doi:10.1101/2021.07.09.451778.

9. Chaumeil PA, Mussig AJ, Hugenholtz P, Parks DH. GTDB-Tk: a toolkit to classify genomes with the Genome Taxonomy Database. Bioinformatics. 2019;36(6):1925–1927. doi:10.1093/bioinformatics/btz848.

10. Eisenhofer R, Weyrich LS. Assessing alignment-based taxonomic classification of ancient microbial DNA. PeerJ. 2019;7:e6594. doi:10.7717/peerj.6594.

11. Zielezinski A, Vinga S, Almeida J, Karlowski WM. Alignment-free sequence comparison: benefits, applications, and tools. Genome Biology. 2017;18(1):1–7. doi:10.1186/s13059-017-1319-7.

12. Segata N, Waldron L, Ballarini A, Narasimhan V, Jousson O, Huttenhower C. Metagenomic microbial community profiling using unique clade-specific marker genes. Nature Methods. 2012;9(8):811–4. doi:10.1038/nmeth.2066.

13. Zhou F, Olman V, Xu Y. Barcodes for genomes and applications. BMC Bioinformatics. 2008;9(1):1–1. doi:10.1186/1471-2105-9-546.

14. Liang Q, Bible PW, Liu Y, Zou B, Wei L. DeepMicrobes: taxonomic classification for metagenomics with deep learning. NAR Genomics and Bioinformatics. 2020;2(1):qaa009. doi:10.1093/nargab/lqaa009.

15. Ounit R, Wanamaker S, Close TJ, Lonardi S. CLARK: fast and accurate classification of metagenomic and genomic sequences using discriminative k-mers. BMC Genomics. 2015;16(1):1–3. doi:10.1186/s12864-015-1419-2.

16. Almeida A, Mitchell AL, Boland M, Forster SC, Gloor GB, Tarkowska A, Lawley TD, Finn RD. A new genomic blueprint of the human gut microbiota. Nature. 2019;568(7753):499–504. doi:10.1038/s41586-019-0965-1.

17. Stewart RD, Auffret MD, Warr A, Wiser AH, Press MO, Langford KW, Liachko I, Snelling TJ, Dewhurst RJ, Walker AW, Roehe R. Assembly of 913 microbial genomes from metagenomic sequencing of the cow rumen. Nature Communications. 2018;9(1):1–1. doi:10.1038/s41467-018-03317-6.

18. Kim D, Song L, Breitwieser FP, Salzberg SL. Centrifuge: rapid and sensitive classification of metagenomic sequences. Genome Research. 2016;26(12):1721–9. doi:10.1101/gr.210641.116.

19. Ounit R, Lonardi S. Higher classification sensitivity of short metagenomic reads with CLARK-S. Bioinformatics. 2016;32(24):3823–5. doi:10.1093/bioinformatics/btw542.

20. Menzel P, Ng KL, Krogh A. Fast and sensitive taxonomic assignment for metagenomics with Kaiju. Nature Communications. 2016;7(1):1–9. doi:10.1038/ncomms11257.

21. Wright RJ, Comeau AM, Langille MG. From defaults to databases: parameter and database choice dramatically impact the performance of metagenomic taxonomic classification tools. bioRxiv. 2022. doi:10.1101/2022.04.27.489753

22. Federhen S. The NCBI taxonomy database. Nucleic Acids Research. 2012;40(D1):D136–43. doi:10.1093/nar/gkr1178.

23. Murray AE, Freudenstein J, Gribaldo S, Hatzenpichler R, Hugenholtz P, Kämpfer P, Konstantinidis KT, Lane CE, Papke RT, Parks DH, Rossello-Mora R. Roadmap for naming uncultivated Archaea and Bacteria. Nature Microbiology. 2020;5(8):987–94. doi:10.1038/s41564-020-0733-x.

24. Huang W, Li L, Myers JR, Marth GT. ART: a next-generation sequencing read simulator. Bioinformatics. 2012;28(4):593–4. doi:0.1093/bioinformatics/btr708.

25. Babbar R, Partalas I, Gaussier E, Amini MR. On flat versus hierarchical classification in large-scale taxonomies. In: Annual Conference on Neural Information Processing Systems; 2013; p. 1824–1832.

26. Randhawa GS, Hill KA, Kari L. ML-DSP: machine learning with digital signal processing for ultrafast, accurate, and scalable genome classification at all taxonomic ranks. BMC Genomics. 2019;20(1):267. doi:10.1186/s12864-019-5571-y.

27. Randhawa GS, Hill KA, Kari L. MLDSP-GUI: an alignment-free standalone tool with an interactive graphical user interface for DNA sequence comparison and analysis. Bioinformatics. 2020;36(7):2258–9.

28. Deschavanne PJ, Giron A, Vilain J, Fagot G, Fertil B. Genomic signature: Characterization and classification of species assessed by chaos game representation of sequences. Molecular Biology and Evolution. 1999;1;16(10):1391–9. doi:10.1093/oxfordjournals.molbev.a026048.

29. Almeida JS, Carrico JA, Maretzek A, Noble PA, Fletcher M. Analysis of genomic sequences by Chaos Game Representation. Bioinformatics. 2001;17(5):429–37. doi:10.1093/bioinformatics/17.5.429.

30. Wang Y, Hill K, Singh S, Kari L. The spectrum of genomic signatures: From dinucleotides to chaos game representation. Gene. 2005;346:173–85. doi:10.1016/j.gene.2004.10.021.

31. Platt J. Probabilistic outputs for support vector machines and comparisons to regularized likelihood methods. Advances in Large Margin Classifiers. 1999;26;10(3):61–74.

32. Arias PM, Alipour F, Hill K, Kari L. DeLUCS - deep learning for unsupervised clustering of DNA sequences. PLoS ONE; 2022; 17(1): e0261531. doi:10.1371/journal.pone.0261531.

33. Gao J, Tan P N. Converting output scores from outlier detection algorithms into probability estimates International Conference on Data Mining (ICDM’06). 2006; p. 212–221. doi:10.1109/ICDM.2006.43

34. Perini L, Vercruyssen V, Davis J. Quantifying the confidence of anomaly detectors in their example-wise predictions. Joint European Conference on Machine Learning and Knowledge Discovery in Databases. 2020; p. 1824–1832.

35. Ramaswamy H, Tewari A, Agarwal S. Convex calibrated surrogates for hierarchical classification. In: Proceedings of Machine Learning Research; 2015; p. 1852–1860.

36. Zheng Z, Cai Y, Li Y. Oversampling method for imbalanced classification. Computing and Informatics. 2015; 34(5): 1017–37.

37. Tax DM. One-class classification: Concept learning in the absence of counter-examples. PhD thesis. Technische Universiteit Delft (The Netherlands). 2002.

