## Supplemental Information for "MT-MAG: Accurate and interpretable machine learning for complete or partial taxonomic assignments of metagenome-assembled genomes"

### S3 Appendix
### Results

In this section, we discuss several details of the computational experiments. In Section 1 we specify the task environment used for running benchmarking task. In Section 2 we discuss the two types of test genomes excluded in the benchmarking comparisons. In Section 3 we introduce the formal definitions of the performance metrics used for benchmarking comparisons. Lastly, we discuss the runtime information for MT-MAG and DeepMicrobes.

#### 1 Task environment

The tasks for MT-MAG were run using python3 and MATLAB R2019b on a x86_64 Ubuntu machine. The tasks for DeepMicrobes were run using python3 on a cluster with RTX6000 partition hosted at the Vector Institute.

#### 2 Excluded test genomes

For the benchmark comparisons between MT-MAG and DeepMicrobes, two types of test genomes were excluded for Task 1 (sparse), as detailed below. First, the ground-truth labels of the test set were determined by running GTDB-Tk [9]. If GTDB-Tk classified the genomes to unnamed species, then these test genomes were excluded as not having ground-truth labels to benchmark the performance of MT-MAG.

Second, the test genomes whose GTDB-Tk-predicted species did not exist in the training set were also excluded. The rationale is that the species in the training set (HGR) form a finite subset of GTDB, and the GTDB-Tk-predicted species for a test genome may not necessarily be in this finite subset. If these genomes would not have been excluded, their MT-MAG and DeepMicrobes classifications could not be correct, since their ground-truth labels had never been seen during training.

For Task 2 (dense), we only excluded the test genomes for which GTDB-Tk-predicted unnamed species.

#### 3 Rigorously defined performance metrics

In the following, we will define the performance metrics for MT-MAG.

Let $G$ denote the test set. Given a test genome $g \in G$ and a taxonomic rank $tr$, let $l(g, tr)$ denote the ground-truth label of $g$ at taxonomic rank $tr$ and let $out^{MT\text{-}MAG}(g, tr)$ denote the label computed by MT-MAG for $g$ at taxonomic rank $tr$. We use "$uc$" to denote uncertain & unattempted classifications at $tr$, that is, if $g$ has an uncertain or unattempted classification at $tr$, then $out^{MT\text{-}MAG}(g, tr)$ is "$uc$."

We denote by $G_c(tr)$ the set of genomes with complete classifications at $tr$ (where $c$ indicates "complete classifications") that is,

$$G_c(tr) = \{g \in G : out^{MT\text{-}MAG}(g, tr) \neq uc\}.$$

We denote by $G'_c(tr)$ the set of genomes with correct classifications down to $tr$, that is,

$$G'_c(tr) = \{g \in G_c(tr) : out^{MT\text{-}MAG}(g, tr) = l(g, tr)\}.$$

We define the *constrained accuracy (of classifying genomes) for taxonomic rank tr* as:

$$CA_g(tr) = \begin{cases} \dfrac{card(G'_c(tr))}{card(G_c(tr))}, & \text{if } card(G_c(tr)) \neq 0 \\ 0, & \text{otherwise} \end{cases}$$

In other words, $CA_g(tr)$ measures how many, out of the test genomes in $G$ with complete classifications at taxonomic rank $tr$, have correct classifications down to $tr$ (See Figure S1).

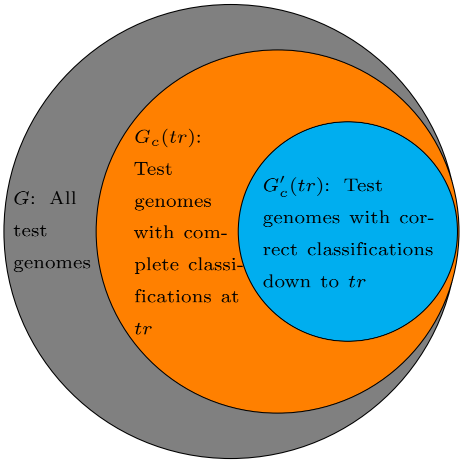


**Fig S1. Composition of the test set for MT-MAG.** A Venn diagram to show the relationship of three types of genomes in $G$ (the set of all test genomes, gray circle) for taxonomic rank $tr$. The set $G_c(tr)$ (orange circle), of the test genomes which have complete classifications at taxonomic rank $tr$, is a subset of $G$, and the set $G'_c(tr)$ (cyan circle), of the test genomes which have correct classifications down to taxonomic rank $tr$, is a subset of $G_c(tr)$. Visually, we have that $CA_g(tr)$ is the ratio of the cyan circle set to the orange circle set, $AA_g(tr)$ is the ratio of the cyan circle set to the gray circle set, and $CR_g(tr)$ is the ratio of the orange circle set to the gray circle set.

We define the *absolute accuracy (of classifying genomes) for taxonomic rank tr* as:

$$AA_g(tr) = \frac{card(G'_c(tr))}{card(G)}.$$

In other words, $AA_g(tr)$ measures how many, out of the test genomes in $G$, have correct classifications down to $tr$ (See Figure S1).

For a given taxonomic rank $tr_j$ in a list of increasingly lower taxonomic ranks $tr_0, tr_1, ..., tr_i$, where $tr_0$ is the root, given a test genome $g \in G$ with a classification path of length $0 \leq h \leq i$, (where $i$ is the maximum length of a classification path), its weight $w(g, tr_j)$ is:

- If the test genome $g$ has a correct classification down to $tr_j$ (i.e., complete classification at $tr_j$, *and* a correct classification at $tr_j$), then the weight is $w(g, tr_j) = 1$.

- If, on the other hand, there exists a $0 < p < j$, such that the test genome $g$ has a correct classification down to $tr_p$, but $g$ does not have a correct classification at $tr_{p+1}$ (the latter condition avoids double counting), then the weight is

$$w(g, tr_j) = max\{0, \frac{1}{j}(p - \frac{1}{2}(min\{h, j\} - p))\}.$$

We define the *weighted accuracy (of classifying genomes) for taxonomic rank* $tr_j$ as

$$WA_g(tr_j) = \frac{\sum_{g \in G} w(g, tr_j)}{card(G)}.$$

We also define the *complete classification rate (for genome classifications) for taxonomic rank tr* as:

$$CR_g(tr) = \frac{card(G_c(tr))}{card(G)}.$$

In other words, $CR_g(tr)$ measures how many, out of the test genomes in $G$, have complete classifications at $tr$ (See Figure S1).

For DeepMicrobes, we define the following performance metrics, that correspond to the MT-MAG metrics $CA_g(tr), AA_g(tr), WA_g(tr)$, and $CR_g(tr)$. To this end, we first define set notations for different categories of test reads for DeepMicrobes. Using these set notations we then formally define the DeepMicrobes performance metrics.

Let $R$ denote the test set of reads. Recall that, given a test read, the output from DeepMicrobes is either a classification of that read at the Species level, or "unclassified." Given a test read $r \in R$, let $l(r)$ denote the ground-truth Species label of $r$, and let $out^{DM}(r)$ denote the label computed by DeepMicrobes for this read if the read was classified, or "*uc*" (unclassified) if the read was not classified.

Denote by $R_c$ the set of classified test reads, that is,

$$R_c = \{r \in R : out^{DM}(r) \neq uc\},$$

where $c$ stands for "classified" (at the Species level).

Denote by $R'_c$ the set of correctly classified reads, that is

$$R'_c = \{r \in R_c : out^{DM}(r) = l(r)\}.$$

We now define the *constrained accuracy (for the classification of reads)*, herein at the Species level, as:

$$CA_r = \begin{cases} \frac{card(R'_c)}{card(R_c)}, & \text{if } card(R_c) \neq 0 \\ 0, & \text{otherwise} \end{cases}$$

In other words, $CA_r$ measures how many, out of the test reads in $R$ with a Species classification, have been correctly classified by DeepMicrobes (See Figure S2).

We define the *absolute accuracy (for the classification of reads)*, herein at the Species level, as:

$$AA_r = \frac{card(R'_c)}{card(R)}.$$

In other words, $AA_r$ measures how many, out of the test reads in $R$, have been correctly classified by DeepMicrobes (See Figure S2). Note that $CA_r$ and $AA_r$ were originally called as $\text{Precision}_{\text{read}}$ and $\text{Recall}_{\text{read}}$ in DeepMicrobes [14].

We define the *weighted accuracy (for the classification of reads)*, herein at the Species level, as being equal to the absolute accuracy $AA_r$. This is because we

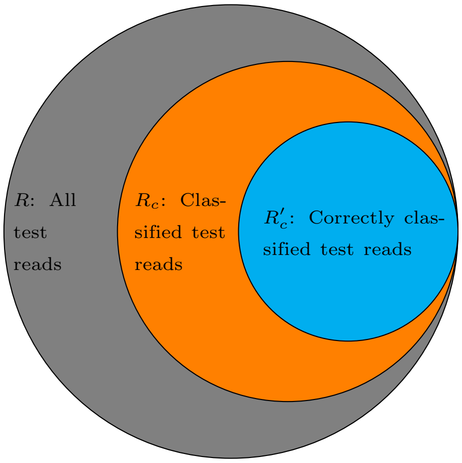


**Fig S2. Composition of the test set for DeepMicrobes.** A Venn diagram to show the relationship of three types of test reads in $R$ (the set of all test reads, gray circle). The set $R_c$ (orange circle), of classified reads, is a subset of $R$, and the set $R'_c$ (cyan circle), of correctly classified test reads, is a subset of $R_c$. Visually, we have that $CA_r$ is the ratio of the cyan circle set to the orange circle set, $AA_r$ is the ratio of the cyan set to the gray set, and $CR_r$ is the ratio of the orange circle set to the gray circle set.

introduced weighted accuracy as a metric meant to combine the classification results of the software for completely classified sequences, with its classification results for partially classified sequences. The latter category does not exist for DeepMicrobes, as it does not provide any classification output about ranks other than Species, hence

$$WA_r = AA_r = \frac{card(R'_c)}{card(R)}.$$

We also define the *classified rate (for the classification of reads)*, herein at the Species level, as:

$$CR_r = \frac{card(R_c)}{card(R)}.$$

In other words, $CR_r$ measures how many, out of the test reads in $R$, could be classified by DeepMicrobes (See Figure S2).

### 4 Runtime

**Table 1.** Summary of MT-MAG and DeepMicrobes's Task 1 and Task 2 runtime in hours. Note that the runtime for DeepMicrobes training is not applicable as we use DeepMicrobes's provided Species-identification model for this task.

| | DeepMicrobes training | MT-MAG training | DeepMicrobes test | MT-MAG test |
|---|---|---|---|---|
| Task 1 | N/A | 121.2 | 24.8 | 61.4 |
| Task 2 | 558.2 | 467.8 | 23.1 | 230.1 |
