## Supplemental Information for "MT-MAG: Accurate and interpretable machine learning for complete or partial taxonomic assignments of metagenome-assembled genomes"

### S2 Appendix
### Methods - MT-MAG algorithm

#### 1 An overview of MLDSP

MLDSP is a software tool introduced in [26], which combines supervised machine learning with digital signal Preprocessing methods into an alignment-free software tool for ultrafast, accurate, and scalable genome classification at all taxonomic ranks. The training set for MLDSP consists of pseudo-concatenated DNA sequences, and the outputs of MLDSP are taxonomic classifications for test or unknown sequences.
MT-MAG uses an enhanced version of MLDSP (eMLDSP) as a subprocess. Note that MLDSP cannot handle the special case where the parent taxon has only one child taxon, and does not output classification confidences. Consequently, MLDSP had to be significantly enhanced to be used as a subprocess of MT-MAG (called eMLDSP).

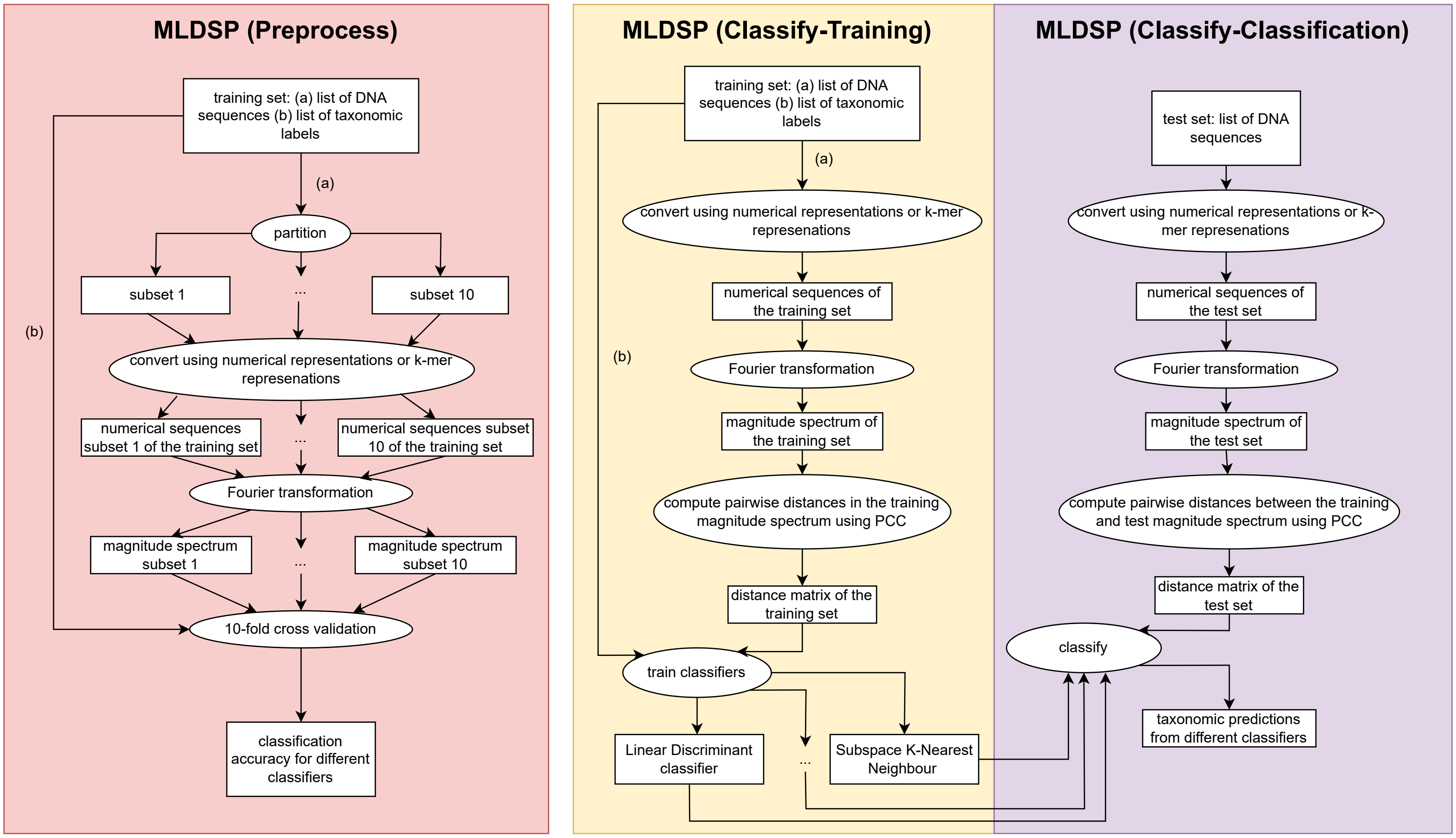


**Fig S1.** An overview of MLDSP, including the main steps to accomplish MLDSP (Preprocessing), MLDSP (Classify-Training) and MLDSP (Classify-Classification). Ellipses represent computation steps. Rectangles represent inputs to and outputs from the computation steps. Note that the training set comprises both (a) DNA sequences and (b) their ground-truth taxonomic labels.

#### 2 Pipeline for single-child classification

Figure S2 illustrates the MT-MAG training phase and classifying phase for classifying two genomes belonging to a given parent taxon, into its only child taxon (single-child classification).

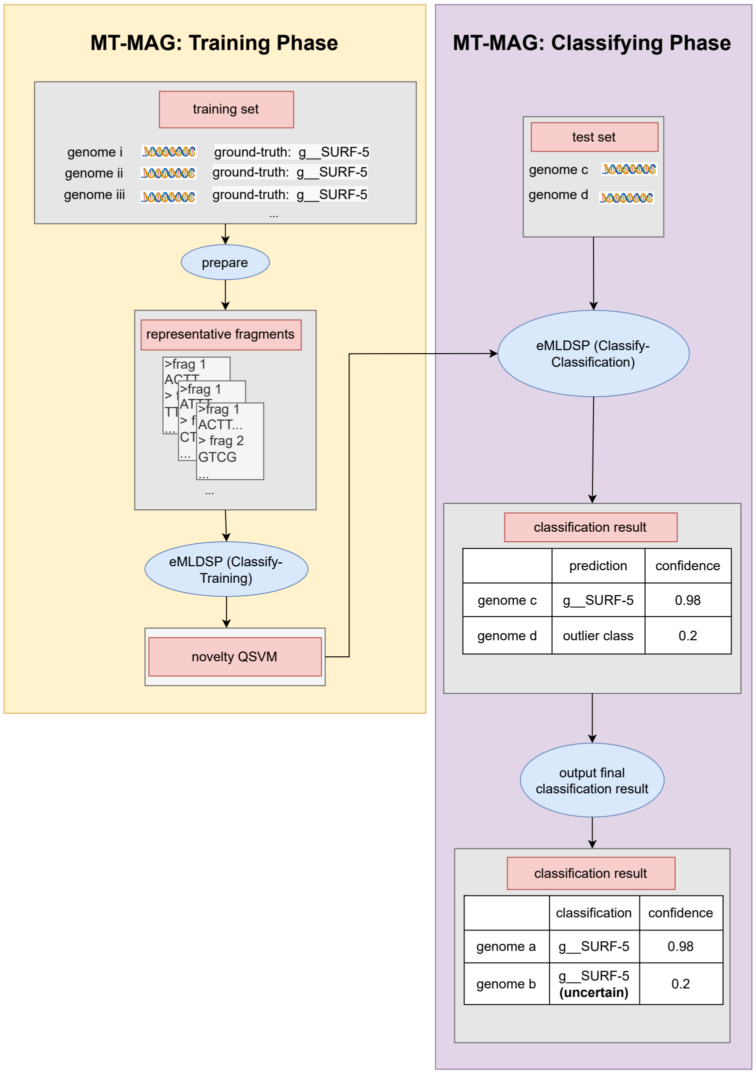


**Fig S2.** MT-MAG pipeline of classifying two genomes, genome $c$ and genome $d$, from the parent taxon Phylum Abyssubacteria into the single-child taxon Class SURF-5 (single-child classification). Blue ellipses represent computation steps. Gray rectangles represent inputs to and outputs from the computation steps. In the training phase, the training set is prepared and given as the input to eMLDSP (Classify-Training), where a novelty QSVM is trained using the entire training set by considering a fraction (default 10%) of the training set to be outliers. In the classifying phase, the test set is given as the input to eMLDSP (Classify-Classification), together with the novelty QSVM from the training phase. eMLDSP (Classify-Classification) outputs a classification and a classification confidence for each genome in the test set. If a genome is classified to be the outlier taxon, then the output is an "uncertain" classification and further classification into children of this child taxon will not be attempted.

#### 3 Training phase

In this section, we will discuss MT-MAG training phase using mathematical notations. First, we give an informal description of the MT-MAG training phase. Second, we give a formal description of eMLDSP, a subprocess for MT-MAG. Third, we give a formal description of the MT-MAG training phase for multi-child classification, with the formal notations defined for eMLDSP. Lastly, we give a detailed description of the MT-MAG training phase for single-child classification.

We start by an informal description of the MT-MAG training phase.

The MT-MAG training phase for the case of multi-child classification consists of two steps: picking stopping thresholds, and training a classifier. A stopping threshold is the result of subtracting a “variability” parameter from the maximum between (a) the minimum of the candidate thresholds (numbers between 0 and 1) that result in a constrained accuracy greater than or equal to a user-specified parameter (default: 90%), and (b) the average of classification confidences of contigs/representative genomic fragments with correct eMLDSP (Preprocessing) classifications.

A stopping threshold for each pair (parent-class, child-class) is stored as a pair (child-class name, child-class stopping threshold), and is utilized as follows. If a test genome is classified to a child-class with a classification confidence that is strictly smaller than the child-class's stopping threshold, then this is considered to be an “uncertain” classification and further classifications of this test genome at lower taxonomic ranks are not attempted. There are three possible cases for an “uncertain” classification. Firstly, the test genome belongs to the uncertain taxon that eMLDSP (Classify-Classification) classification classifies into, however, MT-MAG is not confident about the classification. Secondly, the test genome belongs to another existing taxon. Thirdly, the test genome belongs to a non-existing taxon that is not among the training genomes.

The stopping thresholds were used in the classifying phase to prevent further classification of test data. The classifier was used for classifying test data that have already classified into the parent taxon, into one of the parent taxon's child taxa. For example, in Figure 1, in the training phase of the parent-to-child relationship highlighted in red, the ground-truth labels of all the training genomes should be “rank 2 group 1”, “rank 2 group 2” or “rank 2 group 3”. For the entire taxonomy in Figure 1, MT-MAG trained three classifiers, each corresponding to one of the three parent-to-children relationships. The one from “rank 1 group 2” to “rank 2 group 4”, highlighted in cyan, is for a single-child classification, and the other two (from root to “rank 1 group 1” and “rank 1 group 2”, and from “rank 1 group 1” to “rank 2 group 1”, “rank 2 group 2,” and “rank 2 group 3”) are for multi-child classifications.

The second step of the training phase in the case of multi-child classification is to train a classifier. During this step, for each parent taxon, we trained a QSVM (called fully trained QSVM) using all the training DNA sequences of the parent taxon. This fully trained QSVM will be used, together with the aforementioned stopping thresholds, for the classifying phase.

The MT-MAG training phase for the case of single-child classification does not need a stopping threshold, since the parent taxon has a single child taxon [37]. In this case, all training DNA sequences belong to one class (the single-child taxon), and the goal is for any unknown/test genome to be either categorized as belonging to the present child taxon, or categorized as an outlier taxon (e.g., belonging to a child taxon not represented in the training set). With this goal in mind, a QSVM (called novelty QSVM) is trained, with an optional user-specified outlier fraction (default: 10%) in eMLDSP (Classify-Training). In other words, the novelty QSVM labels a fraction of the training set as outliers (i.e., a second child taxon).

To formally describe the process of picking stopping thresholds in the training phase of MT-MAG (the multi-child classifications case), we now introduce the formal definitions and notations of the concepts involved.

Given a parent-to-child relationship of the multi-child classification type, let $p$ be the parent taxon, let $D_p$ denote the training set of $p$, and let $c$ be a child taxon of $p$, which is a potential classification from eMLDSP (Preprocessing). Let $d$ be a DNA sequence. In our benchmark tasks, $d$ is a contig in the case of Task 1 (sparse), and is a representative genomic fragment in the case of Task 2 (dense).

The subprocess eMLDSP (Preprocessing) for classifying the genomes belonging to a parent taxon $p$ into one of its child taxa can be viewed as a function $M_p(d)$. This function maps each DNA sequence $d$ in the training set (all the genomes from the parent taxon $p$) to a pair, i.e.,

$$M_p(d) = (pred^{M_p}(d),\ \ conf^{M_p}(d))$$

where $pred^{M_p}(d)$ is the taxonomic label of the child taxon of $p$ that was assigned by eMLDSP to the sequence $d$, and the numerical classification confidence $conf^{M_p}(d)$ of this classification.

Using this notation for eMLDSP, the process of determining a stopping threshold $T_p(c)$ for each child taxon $c$ of the parent taxon $p$, can be described as follows.

Denote by $l_p(d)$ the ground-truth label of the child taxon that $d$ belongs to. Note that $l_p(d)$ is a child taxon of $p$.

For a child taxon $c$ of $p$, define $D_p(c)$ to be the set of DNA sequences $d$ in $D_p$ with eMLDSP (Preprocessing) classification being $c$, that is,

$$D_p(c) = \{d \in D_p|\ \ pred^{M_p}(d) = c\}.$$

To determine the stopping threshold for the parent taxon $p$ and its child taxon $c$, we sequentially evaluate a series of candidate thresholds $\alpha \in \{0, 0.01, 0.02, ..., 1\}$, in increasing order of their magnitude. Given a candidate threshold $\alpha$, we denote by $D_p(c, \alpha)$ the set of DNA sequences $d$ in $D_p(c)$ with $conf^{M_p}(d)$ greater than or equal to $\alpha$, that is,

$$D_p(c, \alpha) = \{d \in D_p(c)|\ \ conf^{M_p}(d) \geq \alpha\}.$$

Finally, denote by $D'_p(c, \alpha)$ the set of DNA sequences $d$ in $D_p(c, \alpha)$ whose ground-truth child labels coincide with $c$, that is,

$$D'_p(c, \alpha) = \{d \in D_p(c, \alpha)|\ \ l_p(d) = c\}.$$

We now define the *constrained accuracy* associated to $\alpha$ and $c$ as:

$$CA_p(c, \alpha) = \begin{cases} \dfrac{card(D'_p(c, \alpha))}{card(D_p(c, \alpha))}, & \text{if } card(D_p(c, \alpha)) \neq 0 \\ 1, & \text{otherwise} \end{cases}$$

where $card(S)$ denotes the cardinality of a set $S$, that is, the number of its elements. In other words, $CA_p(c, \alpha)$ measures how many, out of the DNA sequences in $D_p$, with eMLDSP classifications equal to $c$ and classification confidences of their classification greater than or equal to $\alpha$, have been correctly classified by eMLDSP (Preprocessing) (see Figure S3).

The *absolute accuracy* associated to $\alpha$ and $c$ is defined as

$$AA_p(c, \alpha) = \begin{cases} \dfrac{card(D'_p(c, \alpha))}{card(D_p(c))}, & \text{if } card(D_p(c)) \neq 0 \\ 1, & \text{otherwise} \end{cases}.$$

In other words, $AA_p(c, \alpha)$ measures how many, out of the DNA sequences in $D_p$, with eMLDSP classifications equal to $c$, have correct classifications and classification confidences of their classification greater than or equal to $\alpha$ by eMLDSP (Preprocessing) (see Figure S3).

Both $CA_p(c, \alpha)$ and $AA_p(c, \alpha)$ are between 0 and 1, and $CA_p(c, \alpha) \geq AA_p(c, \alpha)$. Indeed, note that $D_p(c, \alpha)$ is a subset of $D_p(c)$, and thus $card(D_p(c, \alpha)) \leq card(D_p(c))$.

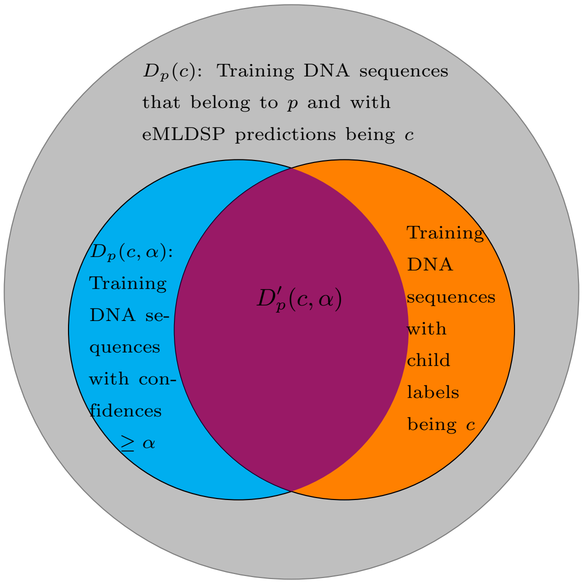


**Fig S3. Training set for MT-MAG (the multi-child classification case).** Relationship among the four types of DNA sequences in $D_p(c)$ (the set of DNA sequences of a parent-taxon $p$ who are predicted by eMLDSP to have child taxon label $c$), for a given candidate threshold $\alpha$. Within the set $D_p(c)$ (gray circle), there are two sets: the set $D_p(c,\alpha)$ (cyan circle), of DNA sequences whose classification confidence as being labelled $c$ is greater than or equal to $\alpha$, and the set of DNA sequences whose ground truth child labels is actually $c$ (orange circle). The intersection of the two sets ($D'_p(c,\alpha)$, violet lens) is the set of DNA sequences $d$ in $D_p(c)$ with classification confidences $\geq \alpha$ and correct eMLDSP (Preprocessing) classifications. Visually, we have that $CA_p(c,\alpha)$ is the ratio of violet lens set to the cyan circle set, and $AA_p(c,\alpha)$ is the ratio of the violet lens set to the gray circle set.

Since the numerators of $CA_p(c,\alpha)$ and $AA_p(c,\alpha)$ are the same, and the denominator of $CA_p(c,\alpha)$ is smaller than or equal to the denominator of $AA_p(c,\alpha)$, it then follows that $CA_p(c,\alpha) \geq AA_p(c,\alpha)$.

The following are the three extreme cases that are possible for $AA_p(c,\alpha)$ and $CA_p(c,\alpha)$:

- When all DNA sequences in $D_p(c)$ have classification confidences strictly less than $\alpha$, we have that both $CA_p(c,\alpha)$ is 1, and $AA_p(c,\alpha)$ is 0. Indeed, in this case, since $D_p(c,\alpha)$ and $D'_p(c,\alpha)$ are empty, it follows that the numerator of $AA_p(c,\alpha)$ are 0, and by definition $CA_p(c,\alpha)$ is 1.
- When all DNA sequences in $D_p(c)$ are correctly classified and have classification confidences greater than or equal to $\alpha$, we have that $CA_p(c,\alpha)$ is 1, and $AA_p(c,\alpha)$ is 1. Indeed, in this case, since $D_p(c) = D_p(c,\alpha) = D'_p(c,\alpha)$, it follows that the numerators and denominators of $CA_p(c,\alpha)$ and $AA_p(c,\alpha)$ are the same.
- When all DNA sequences in $D_p(c)$ are incorrectly classified and have classification confidences greater than or equal to $\alpha$, we have that $CA_p(c,\alpha)$ is 0, and $AA_p(c,\alpha)$ is 0. Indeed, in this case, since $D'_p(c,\alpha)$ is empty, it follows that the numerators of $CA_p(c,\alpha)$ and $AA_p(c,\alpha)$ are both 0.

To intuitively understand what a “reasonably good” $\alpha$ means, we now discuss the case when $CA_p(c,\alpha)$ is 1, but $AA_p(c,\alpha)$ is close to 0. In general, a high $AA_p(c,\alpha)$

indicates a good choice of $\alpha$. If this is not achievable, an indicator of a"reasonably good" choice of $\alpha$ is that $CA_p(c,\alpha)$ is high, while $AA_p(c,\alpha)$ is not too low. Intuitively, the latter requirements mean that *(i)* a high proportion of DNA sequences classified as $c$ with classification confidence $\geq \alpha$, are correctly classified (have ground-truth label $c$) (i.e., high $CA_p(c,\alpha)$), and that *(ii)* in addition, among the training DNA sequences classified to have label $c$, there are sufficiently many training DNA sequences whose classification confidence is at least $\alpha$ (i.e., not too low $AA_p(c,\alpha)$). Note that requiring only that $CA_p(c,\alpha)$ be high could result in situations as follows being considered a "reasonably good" choice for $\alpha$, which would be erroneous: Suppose we have 1,000 training DNA sequences, and that only one DNA sequence has classification confidence $\geq \alpha$ and is correctly classified; then $CA_p(c,\alpha)$ attains the maximum value of 1, but $AA_p(c,\alpha)$ is close to 0.

For each pair comprising a parent taxon $p$ and child taxon $c$, and given a list of candidate thresholds $\{0, 0.01, 0.02, ..., 1\}$, our goal is to determine a stopping threshold $T_p(c)$ from the candidates in this list. The algorithm for computing $T_p(c)$ uses two criteria. Firstly, the algorithm searches for the minimum candidate threshold $\alpha_1 \in \{0, 0.01, 0.02, ..., 1\}$ that results in the constrained accuracy $CA_p(c,\alpha)$ greater than or equal to an optional user-specified constrained accuracy (default: 0.9). Since $AA_p(c,\alpha)$ is a decreasing function of $\alpha$, choosing the minimum threshold candidate $\alpha$ results in the highest possible $AA_p(c,\alpha)$, balancing the twin objectives for $CA_p(c,\alpha)$ and $AA_p(c,\alpha)$. Secondly, the algorithm computes the average $\alpha_2$ of the classification confidences of the training DNA sequences in $D_p(c)$ that are correctly classified as $c$, and computes $max\{\alpha_1, \alpha_2\}$. Furthermore, to account for the additional variability in the test set (which results, in general, in lower classification confidences for the test set compared to the training set), the algorithm accepts an optional user-specified "variability" parameter $v$ between 0 and 1 (default: 0.2). The stopping threshold is now computed as $T_p(c) = max\{\alpha_1, \alpha_2\} - v$.

Algorithm 1 shows the pseudocode for the Stopping Threshold Picking (STP) algorithm. Algorithm 2 shows the pseudocode for the training phase.

**Algorithm 1** Stopping Threshold Picking Algorithm Pseudocode

**Input** $p$ : the parent taxon with more than one child taxon
$CA_u$ : user-specified constrained accuracy (default 0.9)
$v$ : variability (default 0.2)
$\delta$ : the gap between candidate thresholds, default 0.01
**Output** $T_p$: pairs of child taxon of $p$ and its stopping threshold

**procedure** STP$(p, CA_u, v, \delta = 0.01)$
  A $\leftarrow [0, \delta, 2\delta, ..., 1]$ ▷ list of candidate thresholds
  $\alpha_{num} \leftarrow 1/\delta + 1$ ▷ number of candidate thresholds $\alpha$
  $C_p \leftarrow$ child taxa of $p$
  $D_p \leftarrow$ training set of $p$
  **for** $c$ `in` $C_p$ **do**
    $D_p(c) \leftarrow \{d \in D_p : pred^{M_p}(d) = c\}$ ▷ assume non-empty
    $D'_{pc} \leftarrow \alpha_{num}$ of zeros ▷ init, list of $card(D'_p(c, \alpha))$
    $D_{pc} \leftarrow \alpha_{num}$ of zeros ▷ init, list of $card(D_p(c, \alpha))$
    ▷ init, list of confidences for sequences with correct classifications in $D_p(c)$
    $\alpha_{pc} \leftarrow []$
    **for** $d$ `in` $D_p(c)$ **do**
      ▷ the maximum candidate threshold that is smaller than the confidence
      $\alpha_{start} \leftarrow max\{\alpha \leq conf^{M_p}(d) | \alpha \in \mathrm{A}\}$
      $\alpha_{idx} = \frac{\alpha_{start}}{\delta} + 1$ ▷ the index of $\alpha_{start}$ in A
      ▷ if the stopping threshold $\alpha$ is in the first $\alpha_{idx}$ elements of A, $d$ contributes 1 to $card(D_p(c, \alpha))$
      $D_{pc}[1 : d_{idx}] + +$
      **if** $pred^{M_p}(d) = l_p(d)$ **then** ▷ a correct classification
        $D'_{pc}[1 : \alpha_{idx}] + +$ ▷ $d$ contributes 1 to $card(D'_p(c, \alpha))$
        ▷ store the confidence for the correct classification
        append $\alpha_{pc}$ with $conf^{M_p}(d)$
    **for** $card(D'_p(c, \alpha)), card(D_p(c, \alpha)), \alpha$ `in` $D'_{pc}, D_{pc}$, A **do**
      $CA_p(\alpha, c) \leftarrow 1$
      **if** $card(D_p(c, \alpha)) \neq 0$ **then**
        $CA_p(\alpha, c) \leftarrow \frac{card(D'_p(c,\alpha))}{card(D_p(c,\alpha))}$
      **if** $CA_p(\alpha, c) \geq CA_u$ **then** ▷ check the first criterion
        $\alpha_1 \leftarrow \alpha$
        **break** ▷ found the minimum $\alpha \in$ A that satisfies the first criterion
    **if** $card(\alpha_{pc}) \neq 0$ **then** ▷ no correct classifications for child taxon $c$
      $\alpha_2 \leftarrow mean(\alpha_{pc})$ ▷ second criterion
      $T_p(c) \leftarrow max(\alpha_1, \alpha_2) - v$
    Add $(c, T_p(c))$ to $T_p$

### 4 Classifying phase

The classifying phase comprises both *(i)* classifying test genomes with known ground-truth labels, and *(ii)* classifying unknown genomes (without known ground-truth labels). Note that in *(i)*, the ground-truth labels are not used in the classifying phase, and are only needed for computing performance metrics.

In both cases, the classifying phase mimics the hierarchically-structured local classification to classify test/unknown genomes into a leaf taxon (Species-level). The process starts from the root (the highest level parent taxon), and it follows a

**Algorithm 2** Training Phase Pseudocode

**Input** $p$ : the parent taxon with more than one child taxon
$CA_u$ : user-specified constrained accuracy (default 0.9)
$v$ : variability (default 0.2)
**Output** $M_p^{ft}$: a fully trained QSVM, and
$T_p$: pairs of child taxon in $p$ and its corresponding stopping threshold, or
$M_p^{nv}$: a novelty QSVM

```
procedure TRAINING(p, CA_u, v)
    if p has multiple child taxon then                          ▷ multi-child classification
        M_p^{ft} ← a fully trained QSVM trained by the training set of p
        T_p ← STP(p, CA_u, v)
    else                                                        ▷ single-child classification
        M_p^{nv} ← a novelty QSVM trained by the training set of p
```

classification path through increasingly lower taxonomic ranks. This is illustrated in Figure 4, which depicts a fictional hierarchical classification of a genome $g$. The idea of the process is as follows. Suppose that MT-MAG has already determined that the test/unknown genome $g$ belongs to a taxon $p$.

If this is a multi-child classification, denote the fully trained QSVM associated to a parent taxon $p$ by $M_p^{ft}(g)$, where "$ft$" stands for "fully trained." For a input genome $g$, the function $M_p^{ft}(g)$ outputs a pair

$$(pred^{M_p^{ft}}(g),\ \ conf^{M_p^{ft}}(g)),$$

where $pred^{M_p^{ft}}(g)$ is the taxonomic label of the child taxon of $p$ that the fully trained QSVM predicts for $g$, and $conf^{M_p^{ft}}(g)$ is the classification confidence of this classification.

Assume that $pred^{M_p^{ft}}(g) = c_j$, where $c_j$ is one of the child taxa of $p$. Two outcomes are now possible. If the classification confidence for this classification is greater than or equal to the stopping threshold for the pair $p$ and $c_j$, that is, if $conf^{M_p^{ft}}(g) \geq T_p(c_j)$, then MT-MAG outputs "the genome $g$ as belonging to $c_j$, with classification confidence $conf^{M_p^{ft}}(g)$," and then proceeds to further classify $g$ into one of the child taxa of $c_j$. If, on the other hand, the classification confidence is lower than the stopping threshold for this parent-child pair, that is, if $conf^{M_p^{ft}}(g) < T_p(c_j)$, then MT-MAG outputs "classification of $g$ as $c_j$ is uncertain, and the classification confidence is $conf^{M_p^{ft}}(g)$," and stops attempting to classify $g$ further down the taxonomy.

If this is a single-child classification, denote the novelty QSVM associated to a parent taxon $p$ by $M_p^{nv}(g)$, where "$nv$" stands for "novelty." For a input genome $g$, the function $M_p^{nv}(g)$ outputs a pair

$$(pred^{M_p^{nv}}(g),\ \ conf^{M_p^{nv}}(g)),$$

where $pred^{M_p^{nv}}(g)$ is the taxonomic label of the child taxon of $p$ that the novelty QSVM predicts for $g$, and $conf^{M_p^{nv}}(g)$ is the classification confidence of this classification.

Two outcomes are possible. If the novelty QSVM associated to $p$ classifies $g$ as belonging to the single-child taxon $c$ of $p$, that is, if $pred^{M_p^{nv}}(g) = c$, then MT-MAG outputs "the genome $g$ belongs to $c$, with classification confidence $conf^{M_p^{nv}}(g)$," and then proceeds to further classify $g$ into one of the child taxa of $c$. If, on the other hand, if $M_p^{nv}$ classifies $g$ to the outlier taxon, that is, if $M_p^{nv} \neq c$, then MT-MAG outputs "classification of $g$ as $c$ is uncertain, and the classification confidence is $conf^{M_p^{nv}}(g)$," and stops attempting to classify $g$ further down the taxonomy.

Algorithm 3 shows the pseudocode for the classifying phase.

**Algorithm 3** Classifying Phase Pseudocode

**Input** $g$ : the test/unknown genome to be classified
**Output** $cp$ : the classification path with classification confidences for $g$

**procedure** CLASSIFYING($g$)
  $cp \leftarrow (\text{root}, 1)$ ▷ init, classification path with classification confidences
  $t \leftarrow \text{root}$ ▷ init, current taxon
  **while** $t$ is not a leaf taxon **do**
    **if** $t$ has more than one child taxon **then** ▷ multi-child classification
      **if** $conf^{M_p^{ft}}(g) \geq T_p(t)$ **then** ▷ confidence passes stopping threshold
        $t \leftarrow pred^{M_p^{ft}}(g)$
        append $cp$ with $(t, conf^{M_p^{ft}}(g))$
      **else** ▷ confidence does not pass stopping threshold
        append $cp$ with ($t$, "uncertain", $conf^{M_p^{ft}}(g)$)
        **break**
    **else** ▷ single-child classification
      $sc \leftarrow$ the single child taxon of $t$
      **if** $pred^{M_p^{nv}}(g) = sc$ **then** ▷ classified to the single child taxon
        $t \leftarrow sc$
        append $cp$ with $(sc, conf^{M_p^{nv}}(g))$
      **else** ▷ classified to the outlier taxon
        append $cp$ with ($sc$, "uncertain", $conf^{M_p^{nv}}(g)$)
        **break**

### 5 Optimizing MT-MAG

A careful analysis of MT-MAG's time complexity reveals that a significant part of its runtime comes from its training phase. In addition, in the task of classification of a test/unknown genome, not all novelty QSVMs, fully trained QSVMs, and stopping thresholds computed during the training phase are used in the classifying phase. Indeed, for a test/unknown genome, only the novelty/fully trained QSVM's and stopping thresholds local to its classification path will be actually used for the classification.

Thus, MT-MAG can be optimized to prevent computation of unnecessary novelty/fully trained QSVMs, and unnecessary stopping thresholds, as follows. First, with the exception of the root taxon, MT-MAG will only train a novelty/fully trained QSVM of a parent taxon $p$ if there are test/unknown genomes that have been classified to $p$ by the novelty/fully trained QSVM of the parent of $p$. Second, the algorithm for determining the stopping thresholds can be optimized by computing only the stopping thresholds for pairs of parent taxon $p$ and child taxon $c$, in the case where *(i)* there are test/unknown genomes that have been classified to $p$ by the novelty/fully trained QSVM of the parent of $p$, and *(ii)* there are test/unknown genomes that have been classified to $c$ by the fully trained QSVM of $p$.
