## Supplemental Information for "MT-MAG: Accurate and interpretable machine learning for complete or partial taxonomic assignments of metagenome-assembled genomes"

### S1 Appendix
### Materials - Datasets and task description

#### 1 Genome size, contig count, percent GC distributions

The following histograms show the genome size, contig count and percent GC distributions for different datasets in Task 1 (sparse) and Task 2 (dense). Specifically, we have:

- Figure S1 – Task 1 (sparse) training set: unclassified MAGs, and genomes from human-specific reference (HGR) database.
- Figure S2 – Task 1 (sparse) test set: high-quality MAGs reconstructed from human gut microbiomes from a European Nucleotide Archive study titled "A new **g**enomic **b**lueprint of the **h**uman **g**ut **m**icrobiota" (GBHGM) [16].
- Figure S3 – Task 2 (dense) training set: genomes from Genome Taxonomy Database (GTDB) R06-RS202.
- Figure S4 – Task 2 (dense) test set: microbial genomes from metagenomic sequencing of cow rumen, which were derived from 43 Scottish cattle [17].

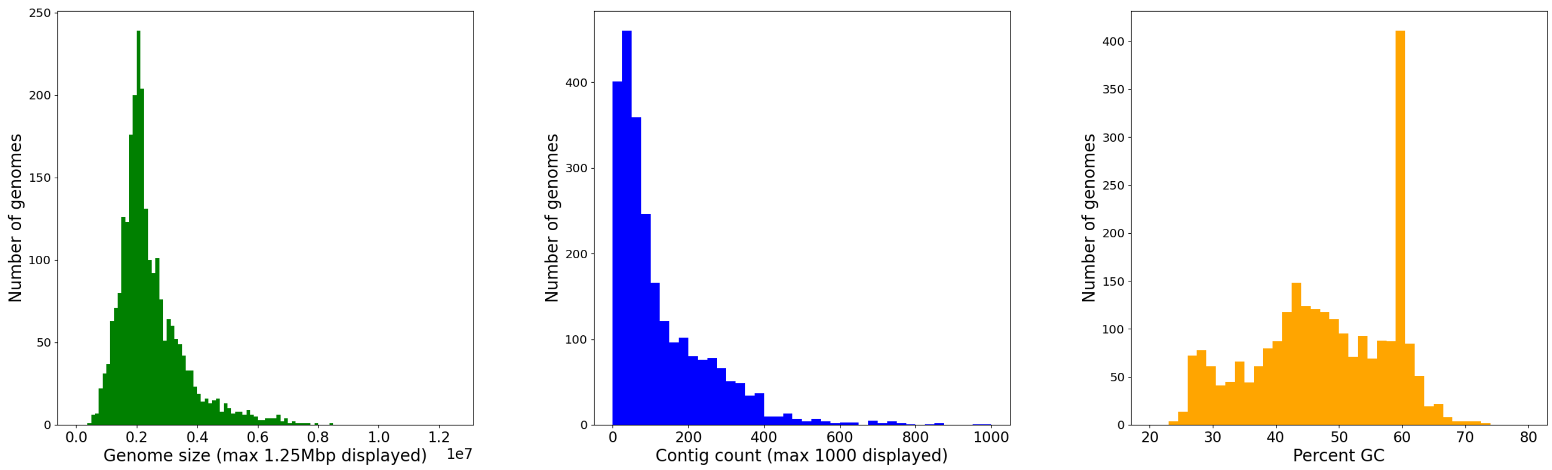


**Fig S1.** Genome size, contig count and percent GC distributions for all genomes in the HGR database. Recall that HGR comprises 1,952 MAGs and 553 microbial gut Species-level genome representatives from the human-specific reference database. From the histograms, we observe that the genome sizes are centered around 2 Mbp; the contig count is right-skewed and peaks at around 38; the percent GC peaks at around 59%.

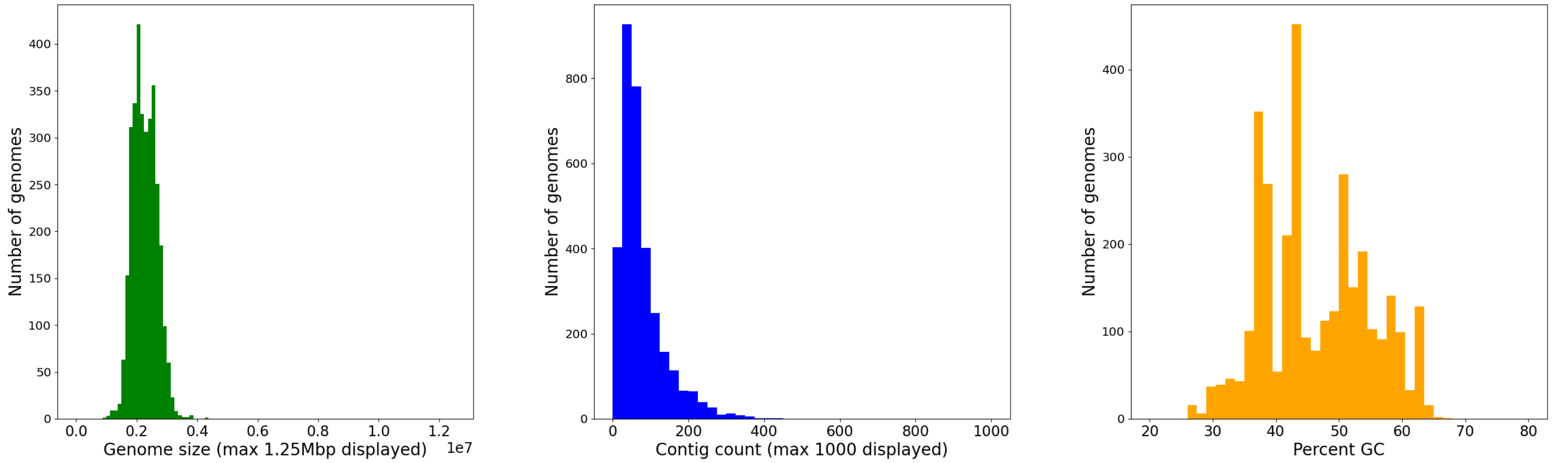


**Fig S2.** Genome size, contig count and percent GC distributions for all GBHGM MAGs. Recall that GBHGM comprises 3,269 high-quality MAGs reconstructed from human gut microbiomes from a European Nucleotide Archive study titled "A new **g**enomic **b**lueprint of the **h**uman **g**ut **m**icrobiota" (GBHGM) [16]. From the histograms we observe that the genome sizes are centered around 2 Mbp; the contig count is right-skewed and peaks at around 38; the percent GC peaks at around 44%.

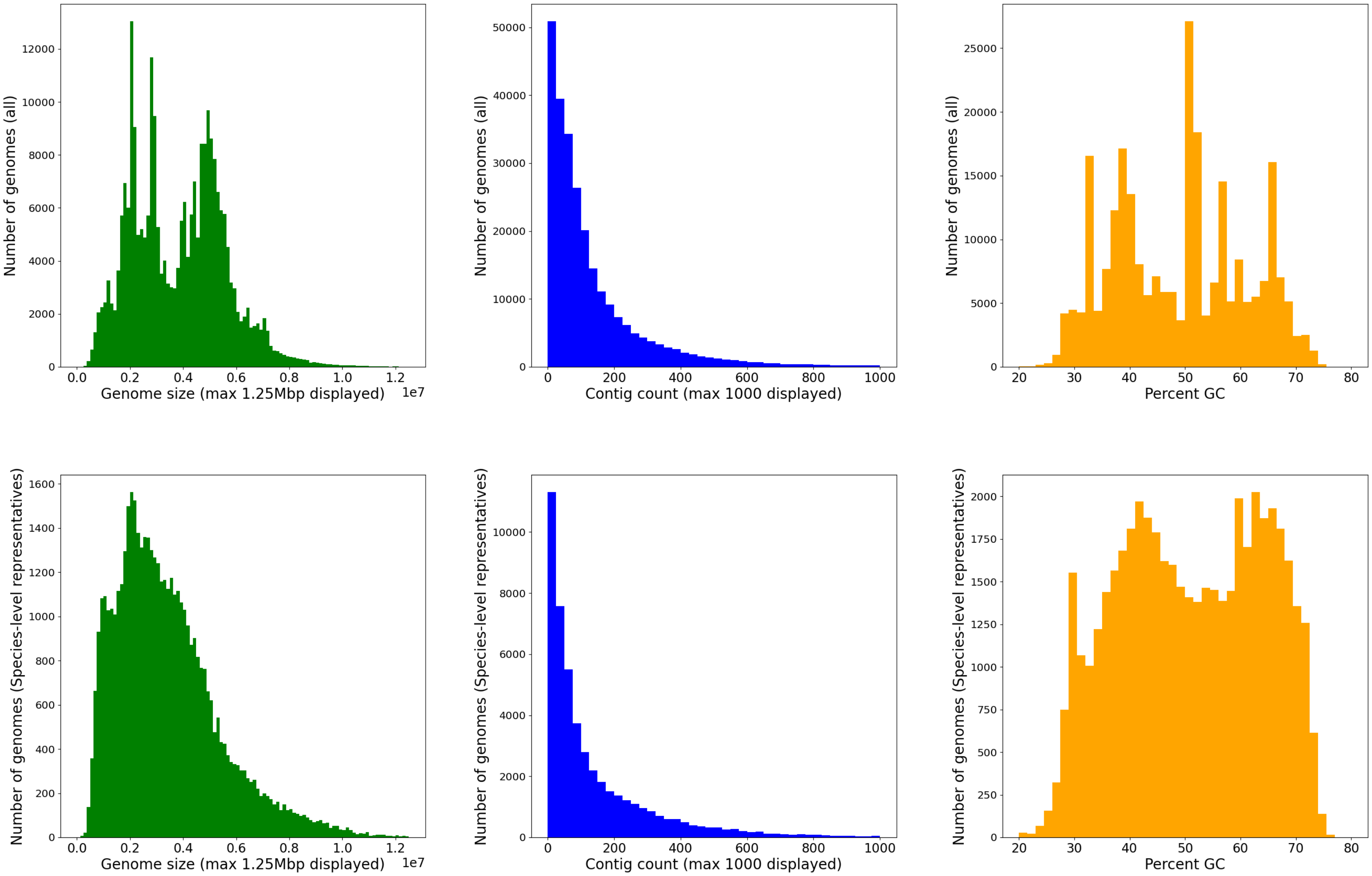


**Fig S3.** Genome size, contig count and percent GC distributions for all genomes (top panels) and Species-level representative genomes in GTDB (bottom panels). Recall that the full GTDB comprises 311,480 bacteria genomes and 6,062 archaeal genomes. From the histograms in the top panels, generated for all genomes in GTDB, we observe that the genome sizes are multimodal and peak at around 2 Mbp, 3 Mbp and 5 Mbp; the contig count is right-skewed and peaks at around 13; the percent GC peaks at around 51%. From the histograms in the bottom panels, generated for Species level representatives in GTDB, the genome sizes are right-skewed and peak at around 2 Mbp; the contig count is right-skewed and peaks at around 13; the percent GC peaks at around 41% and 64%.

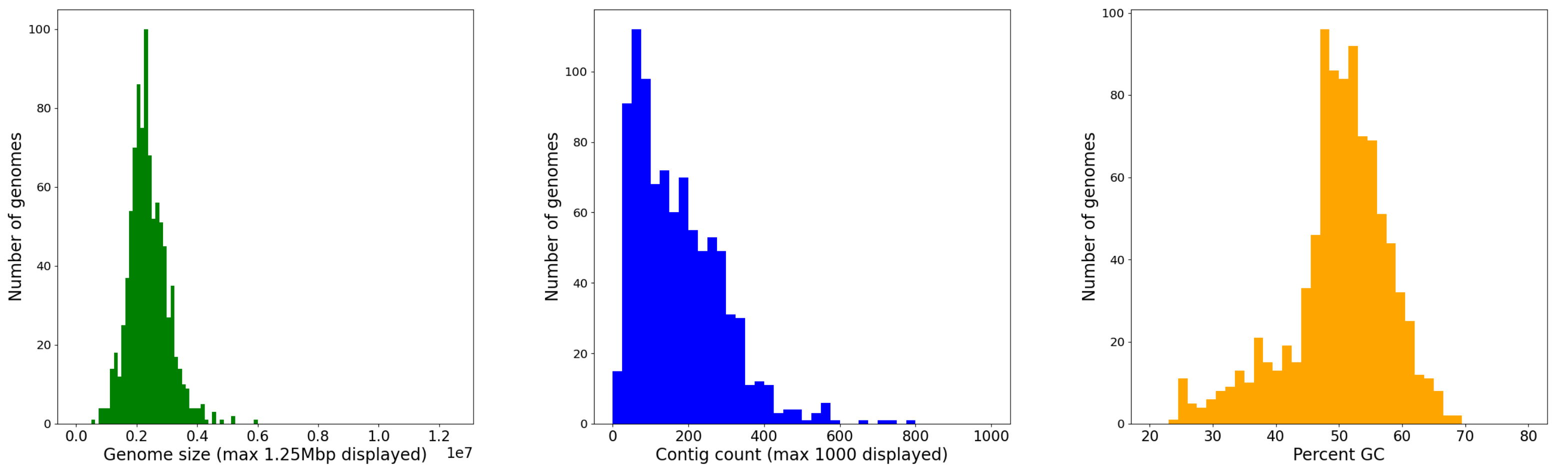


**Fig S4.** Genome size, contig count and percent GC distributions for all 913 cow rumen MAGs. Recall that there are 913 "draft" bacterial and archaeal genomes assembled from rumen metagenomic sequence data derived from 43 Scottish cattle. From the histograms, we observe that the genome sizes are centered around 2 Mbp; the contig count is right-skewed and peaks at around 63; the percent GC peaks at around 50%.

#### 2 Seed fixing

To ensure our results are replicable, our presented results were produced by setting the random seed to 0 for all processes that involve randomness (unless otherwise mentioned).

#### 3 Special cases

- *Dataset being too large for Task 1 (sparse).* For the Phylum level classification, as well as for the over-represented lineages from Class to Genus level classifications, we randomly selected 10% of contigs from any child taxon with more than 100 contigs. This reduced sampling bias in the datasets and reduced computational complexity without omitting any taxon within the taxonomy.

- *Dataset being too small for Task 2 (dense).* One type of special case concerns the taxa with insufficient number of representative genomic fragments. More specifically, to perform five-fold cross-validation, each child taxon needs to have at least five representative genomic fragments. To address this issue, in such cases, and for the Phylum to Genus level taxa, the child taxa with fewer than five representative genomic fragments were removed.

- *Dataset being too large for Task 2 (dense).* For Domain, Phylum, and over-represented lineages from Class to Genus, where eMLDSP consumes an extreme large amount of random access memory, we randomly selected 10% of the representative genomes from any child taxon with more than 100 representative genomes. For less-well represented lineages from Classes to Genera, we selected all representative genomes. For Species, we selected all representative and non-representative genomes.

- *Imbalanced classification for Task 1 (sparse) and Task 2 (dense).* One scenario which could lead to problematic classifications is that of imbalanced taxon sizes.

This is because significant differences in child taxon sizes may violate the assumption, necessary for most classification algorithms, that the number of the training instances for each class be roughly the same. Imbalanced class sizes may pose a challenge for predictive performance, especially for the classes with few training instances [36]. In general, there is a trade-off between balanced class sizes and the amount of variability reflected in each class. In the computational experiments in this paper, the situation of imbalanced taxon sizes was dealt with differently, depending on the taxonomic rank. For high-level classifications (i.e., Domain to Phylum, Phylum to Class, Class to Order, Order to Family), no pruning of over-sampled child taxa was performed. This is because the number of (representative) genomes in a child taxon is proportional to the amount of variability in the taxon and, for high-level classifications, the differences in the amounts of variability of different child taxa can be significant. Since the training instances are intended to capture and represent the differences in the amount of variability, the oversized child taxa must be preserved as being reflective of their respective amounts of variability, and were not pruned.

The situation is different for low-level classifications (i.e., Family to Genus, and Genus to Species), where the differences in the amounts of variability of various child taxa is much smaller, and imbalanced taxon sizes can have a negative impact on the training model and its performance. In these latter cases, all training child taxa were pruned to relatively similar sizes as follows. After sampling, the number of contigs/representative genomic fragments in each child taxon was counted. If the number of contigs/representative genomic fragments was greater than 30, then 30 contigs/representative genomic fragments were randomly selected and used for training.

#### 4 DeepMicrobes training

For Task 2 (dense), via a hyper-parameter search, we found the attention model with the following hyper-parameter values yielded the best performance: the embedding dimension being 50, the batch size being 1024, the learning rate starting with 0.001. 30,000 150 basepair (bp) paired-end reads were simulated from HiSeq 2500 error model in ART Illumina read simulator [24], with the mean fragment size of 200, standard deviation of 50 bp per species and seeds 1, 2 and 11.
